# Distinct effects of phyllosphere and rhizosphere microbes on invader *Ageratina adenophora* during its early life stages

**DOI:** 10.1101/2024.01.09.574925

**Authors:** Zhao-Ying Zeng, Jun-Rong Huang, Zi-Qing Liu, Ai-Ling Yang, Yu-Xuan Li, Yong-Lan Wang, Han-Bo Zhang

**Affiliations:** State Key Laboratory for Conservation and Utilization of Bio-Resources in Yunnan, Yunnan University, Kunming, China; School of Ecology and Environmental Science, Yunnan University, Kunming, China

## Abstract

Microbes strongly affect invasive plant growth. However, how the phyllosphere and rhizosphere soil microbes distinctively affect seedling mortality and the growth of invasive plants across ontogeny under varying soil nutrient levels remains unclear. In this study, we used the invader *Ageratina adenophora* to evaluate these effects in plant growth chambers. We found that leaf litter harboured more potential pathogens and thus had more adverse effects on seed germination and seedling survival than soil inoculation. Microbial inoculation at different growth stages altered the microbial community and microbial functions of seedlings, and earlier inoculation had a more adverse effect on seedling survival and growth. In most cases, the soil nutrient level did not affect microbe-mediated seedling growth and the relative abundance of the microbial community and functions involved in seedling growth. The effects of some microbial genera on seedling survival are distinct from those on growth. Moreover, the *A. adenophora* seedling-killing effects of fungal strains isolated from dead seedlings by nonsterile leaf inoculation litter exhibited significant phylogenetic signals, by which strains of *Allophoma* and *Alternaria* generally caused high seedling mortality. Our study stresses the essential role of *A. adenophora* litter microbes in population establishment by regulating seedling density and growth.

## Introduction

Plant-modified soil properties affect the performance of plants, which are termed ‘plant–soil feedbacks (PSFs)’. PSFs can affect species coexistence (Bever et al., 1997; van der Putten et al., 2013) and local plant community composition and dynamics (Bauer et al., 2017; Kardol et al., 2007; Teste et al., 2017). For plant invasion, PSFs are usually positive because of escaping soil pathogens and recruiting some beneficial microbes (Liu et al., 2023; Mitchell & Power, 2003; Xu et al., 2012) or negative effects because of accumulating local pathogens (Callaway et al., 2013; Flory & Clay, 2013; Zhang et al., 2020).

Similar to soil, leaf litter can also affect plant growth, species diversity and community structure (Liu et al., 2017; Ma et al., 2020; Olson & Wallander, 2002), thus playing important roles in population establishment and community dynamics (Jessen et al., 2023; Lamb, 2008; Xiong & Nilsson, 1999). However, related research has focused mainly on physical (e.g., maintaining soil moisture and temperature, increasing nutrition and reducing light) or chemical effects (e.g., releasing allelochemicals) (Demey et al., 2013; Jessen et al., 2023; Möhler et al., 2018; Zhang et al., 2017) but has rarely focused on leaf microbial effects. Until 2017, Whitaker et al. (2017) extended the PSF to aboveground tissues (including leaf, stem and floral tissues), termed “plant‒ phyllosphere feedbacks (PPFs)”, and found that all four Asteraceae species experienced stronger negative PPFs than PSFs. Subsequently, this team further verified that all ten tested Asteraceae plants experienced negative PPFs (Zaret et al., 2021). The lack of strong mutualists and relatively high abundance of pathogens in the phyllosphere may account for the negative PPFs.

In addition to microbial sources (i.e., soil vs leaf litter), ontogeny (seedling growth stage) and soil nutrient levels can affect plant‒microbe interactions. For example, seedlings showed distinct sensitivity during the growth stage, and younger seedlings were more susceptible to infection by soil microbes because of fewer defense resources (Geisen et al., 2021; Jevon et al., 2020). Interestingly, leaf litter has an adverse effect on seedling emergence but a positive effect on later plant growth (Abbas et al., 2023; Möhler et al., 2021; Zhang et al., 2022); litter also has a stronger negative effect on earlier vegetation growth than on the elder (Loydi et al., 2013; Wang et al., 2022; Xiong & Nilsson, 1999). Moreover, plants enrich distinct microbes under different nutrient conditions and affect plant performance (Dostál, 2021; Gustafson & Casper, 2004; Widdig et al., 2020). For example, the bacterial diversity in duckweed plants was reduced under nutrient‒deficient conditions, but the abundance of Firmicutes increased (Bunyoo et al., 2022), and members of Firmicutes have been reported to promote plant stress tolerance (Xu et al., 2018). Ai et al. (2018) reported that nutrient additions cause crops to enrich some bacteria and fungi from soil and increase yield.

*Ageratina adenophora* (Sprengel) R.M. King & H. Robinson (Asteraceae), known as Crofton weed or Mexican devil weed, has invaded more than 30 countries and areas, including South Africa, Australia, New Zealand, Hawaii, India and China (Gu et al., 2021; Julien & Griffiths, 1998). It is a perennial weed and can produce high yields of seeds with a high germination rate (Lu et al., 2006; Parsons WT, 1992). This weed commonly grows in monocultures, but a high density of seedlings is not common in the wild. Previous studies have shown that *A. adenophora* can enrich the beneficial soil microbial community to facilitate invasion (Niu et al., 2007; Zhao et al., 2019); in contrast, *A. adenophora* leaves harbor diverse fungal pathogens that can cause adverse effects on itself seed germination and growth (Kai Fang et al., 2019; K. Fang et al., 2021). Thus, it is interesting to determine whether leaf microbes play a distinct role from soil microbes in regulating *A. adenophora* seedling density and whether these effects change with the *A. adenophora* growth stage and soil nutrition level.

In this study, we inoculated *A. adenophora* with soil or leaf litter at three stages, 0 d, 21 d, and 28 d after sowing, and transplanted seedlings to grow in soils with high or low nutritional levels. We first determined the germination, seedling survival and growth of the *A. adenophora* plants. Then, we characterized the bacterial and fungal communities of the soils and leaf litter as inoculation sources, as well as the microbial communities enriched in the leaves and roots of the *A. adenophora* seedlings after growing; we also isolated the fungi associated with the dead seedlings and tested their seedling-killing effects on *A. adenophora*. Finally, we correlated the microbial community with *A. adenophora* seedling mortality and growth.

We hypothesized that the microbial communities associated with leaf litter and rhizosphere soils can account for the differential effects on *A. adenophora* seedling mortality and growth during different growth stages when growing under different nutrient conditions. We expected that 1) leaf litter would have more adverse effects on seed germination, seedling survival and growth than soil, as leaf litter often harbours more plant pathogens, and 2) inoculation at different growth stages would change the microbial community enriched by seedlings and thus affect seedling growth. Earlier inoculation had a greater adverse impact on seedling growth than later inoculation, as younger seedlings were more sensitive to pathogen infection than older seedlings. 3) The nutrient level influences seedlings to recruit microbes and thus affects seedling growth.

## Results

### Effects of leaf litter and rhizosphere soil on the mortality and growth of *A. adenophora* seedlings

At the G0 timepoint, sterile leaf inoculation significantly delayed germination time more than did soil and sterile leaf inoculation, as well as the control (nothing inoculated) (Fig. 1a, *P* < 0.05). Leaf and soil inoculation had no distinct effects on the germination rate (Fig. 1b, *P* > 0.05). In addition, the inoculation of sterile and non–sterile leaves at G0 caused a high mortality rate (19.7% vs 96.7%) for seedlings growing in petri dishes (Fig. 1c, Fig. S1). Only nonsterile leaves caused a low percentage of seedling death (8.4%) when the seedlings were inoculated at G21 (Fig. 1c). Two weeks after transplanting these seedlings into the cups, leaf inoculation caused significantly greater seedling mortality than did soil inoculation (*P* < 0.001); the nonsterile sample caused greater seedling mortality than did the sterile sample, especially leaf inoculation during the G0 and G21 periods. Moreover, nonsterile leaf inoculation at earlier stages significantly increased seedling mortality compared with that at later stages (Fig. 1d, *P* < 0.05). However, seedling mortality did not differ between the high- and low-nutrient conditions, regardless of leaf or soil inoculation (Fig. 1d, both *P* > 0.05).

**Figure 1.**
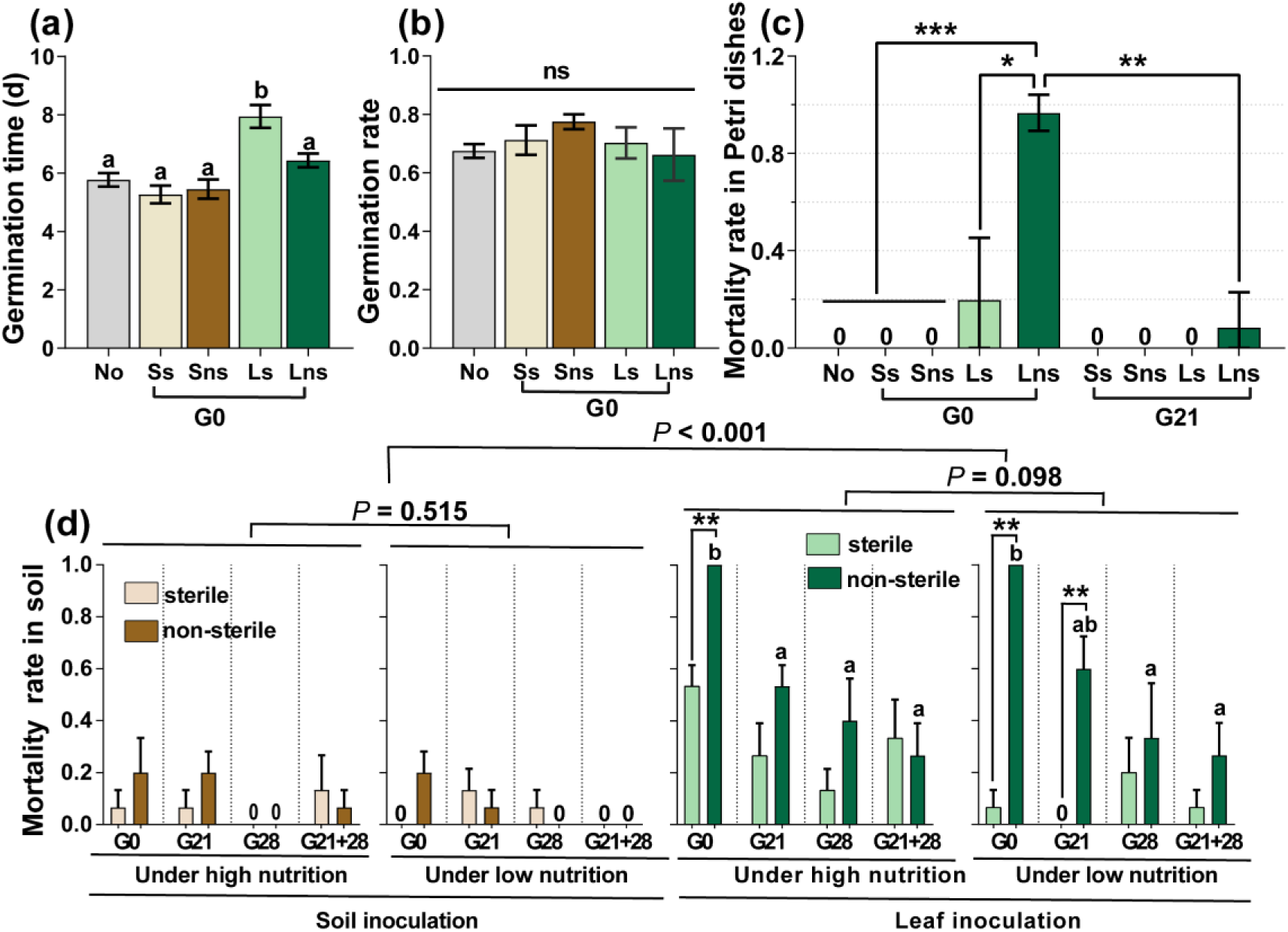
*A. adenophora* seed germination and seedling mortality. Seed germination time and rate after soil or leaf inoculation at the G0 (a-b) and seedling mortality rate after soil or leaf inoculation at the G0 or G21 and subsequent growth in Petri dishes (c). Seedling mortality rate after two weeks when seedlings were transplanted into soil (d). No: nothing inoculated, Ss: sterile soil; Sns: non–sterile soil; Ls: sterile leaf; Lns: non–sterile leaf; G0, inoculated on the day of germination; G21, inoculated on the 21st day after germination; G28, inoculated on the 28th day after germination; G21+28, inoculated on both the 21st and 28th days after germination. * *P* < 0.05, *** *P* < 0.001. Error bars depict the standard error. Different lowercase letters represent significant differences among the different inoculation time treatments (*P* < 0.05). No lowercase letters indicate no differences among the four inoculation time treatments under the same nutrient level (*P* > 0.05).

With the exception of nutrient level, inoculation source and time, as well as their interaction with nutrient level, significantly affected the microbial role in the total dry biomass of *A. adenophora* (*P* < 0.05, Fig. 2a). Soil and leaf microbial effects on seedling biomass interacted with soil nutrient level and inoculation time: when inoculated at the G0 timepoint, both soil and leaf microbes had adverse effects on *A. adenophora* growth, as all seedlings inoculated with nonsterile leaves died, and those inoculated with nonsterile soil grew poorer than those inoculated with sterile soil under both low- and high-nutrition conditions; when inoculated at the G21 timepoint, both soil and leaf microbes had significantly positive effects only under high-nutrition conditions; when inoculated at the G28 timepoint, only soil microbes promoted seedling growth under high-nutrition conditions; and when inoculated at the G21+28 timepoint, only leaf microbes had a significantly positive effect on seedling growth under low-nutrition conditions (Fig. 2b).

**Figure 2.**
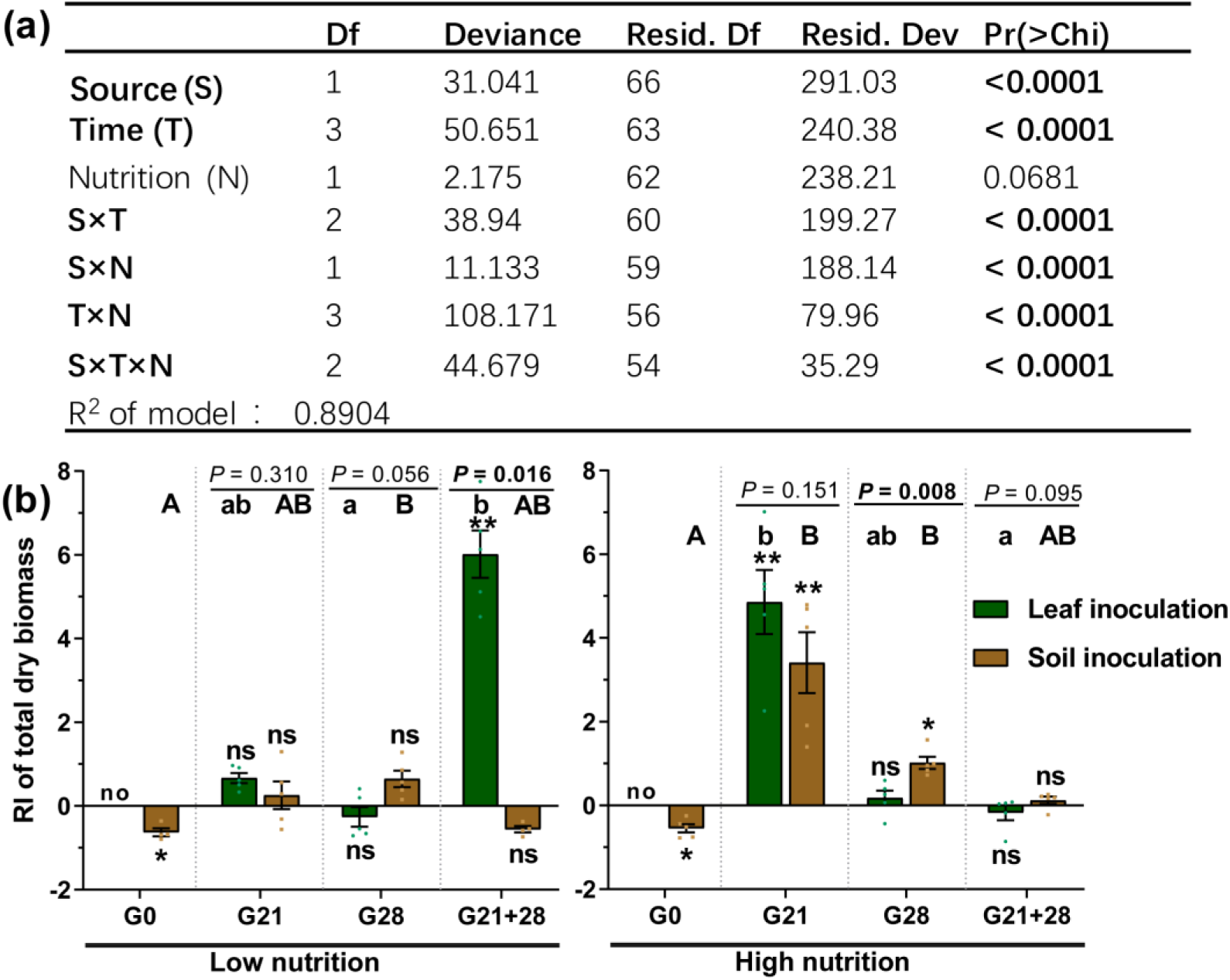
GLM analysis (a) and comparison (b) of the role of microbes in seedling growth based on total dry biomass (RI) between leaf and soil inoculated at low and high nutrient levels across four inoculation time points. Source refers to the leaf or soil inoculation source, time refers to the four inoculation time treatments, and nutrition refers to the nutrient level of the soil. For G0, G21, G28 and G21+28, see Figure 1. Error bars depict the standard error. The difference in the RI among the different inoculation time treatments is indicated by different capital and lowercase letters for the soil and leaf inoculation treatments, respectively. The asterisks indicate that the RIs are significantly different from zero. “No” indicates no surviving seedlings at harvest. *P* < 0.05 is shown in bold.

### Correlations of the microbial community and potential functions of inocula with *A. adenophora* seedling mortality at the early stage

The soil and leaf inocula had distinct microbial diversity, community compositions and potential functions. The microbial diversity and richness were greater in the soil than in the leaf litter (Fig. S2). Top three core bacteria were *Rhodoplanes* (5.65%), *Bradyrizhobium* (4.80%) and some unclassified Alphaproteobacteria (4.56%) for soil and *Pseudomonas* (30.33%), *Massilia* (17.45%) and *Sphingomonas* (16.35%) for leaf (Fig. 3ab). Top three core fungal genera were *Mortierella* (7.00%), *Inocybe* (5.36%) and *Neonectria* (3.63%) for soil and *Didymella* (27.30%) *Alternaria* (8.95%) and *Cryptococcus* (4.44%) for leaf (Fig. 3ef). Plant bacterial pathogens (2.29%) were the potential function of the core bacterial taxa in leaves but not in soil; soil had a greater abundance of nitrogen circle–related function (50.98%) than did leaves (17.55%) (Fig. 3cd, Table S1-2). The abundance of plant fungal pathogens was greater in leaves (68.20%) than in soil (33.93%) (Fig. 3gh, Table S3-4).

We further correlated the top 30 microbial genera of leaf and soil inocula with seed germination and seedling mortality in response to inoculation with nonsterile inocula at G0. The abundances of both the soil and leaf microbial genera were related to the seedling mortality rate (MR) but not to the germination time (GT) or germination rate (GR) (Fig. 3ij). Specifically, the leaf core bacterial genera *Pseudomonas*, *Sphingomonas*, *Massilia*, *Variovorax*, *Aureimonas*, and *Agrobacterium* were positively correlated with MR, but the soil core bacteria, including Alphaproteobacteria_unclassified, *Rhodoplanes*, Vicinamibacteraceae_unclassified, and *Pedomicrobium*, were negatively correlated with MR (Fig. 3abi). The leaf core fungal genera *Didymella*, Pleosporales_unclassified, *Subplenodomus*, and *Bulleromyces* were positively correlated with MR, but the soil core fungal genera *Mortierella*, *Hypocrea*, *Pochonia* and *Volutella* were negatively correlated with MR (Fig. 3efj).

**Figure 3.**
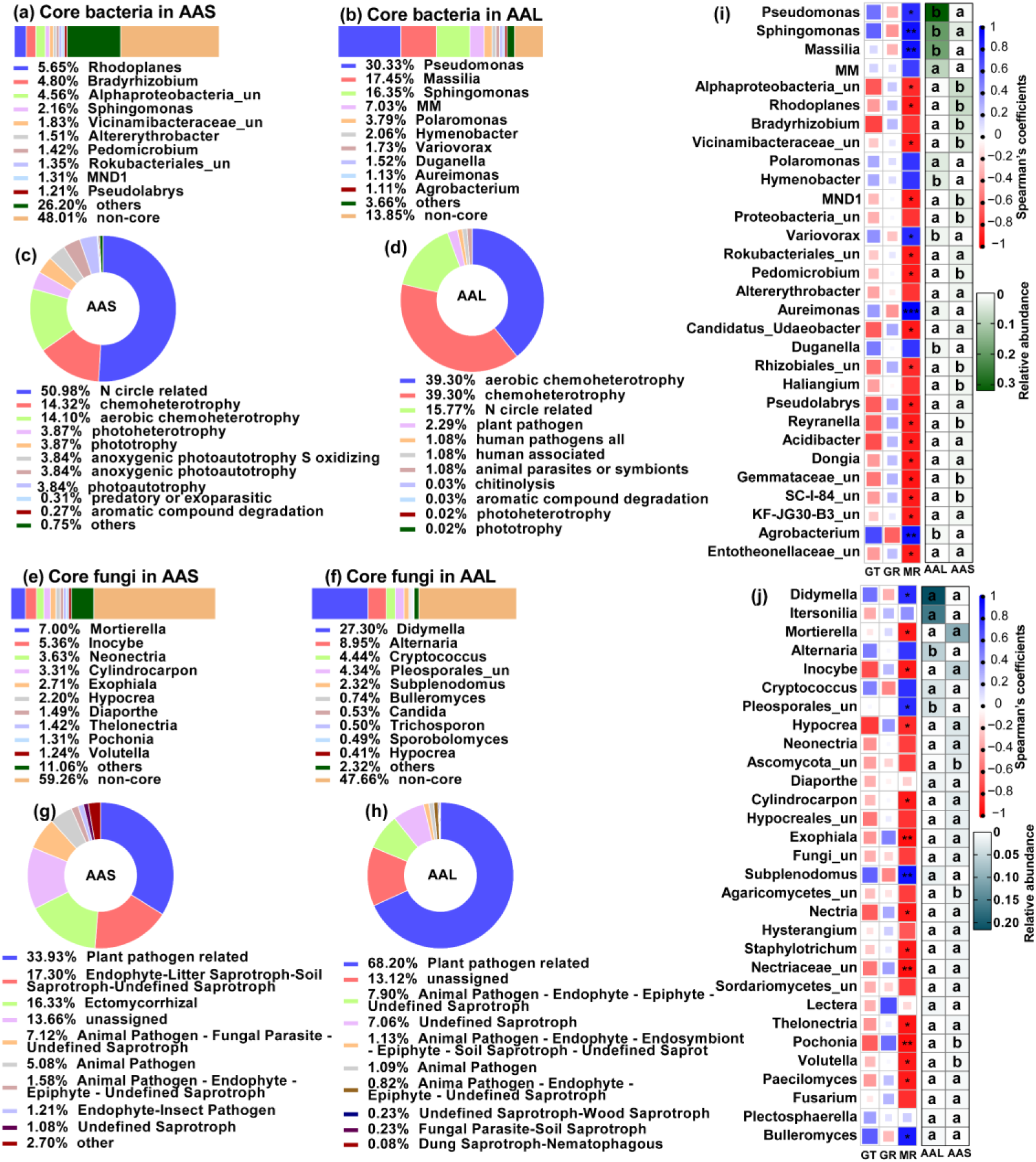
Microbial community composition and potential functional differences between leaf litter (AAL) and rhizosphere soil (AAS) inocula (a-h), as well as correlations of microbial genera with seed germination and seedling mortality (i-j). Core bacteria (a-b) and fungi (e-f) in the AAS and AAL groups. The potential functions of the core bacteria (c-d) and fungi (g-h) in the AAS and AAL. Correlations of the relative abundance of the top 30 bacterial (i) and fungal (j) genera with germination time (GT), germination rate (GR) and mortality rate (MR). Only the top 10 core taxa and potential functional groups are shown in the figures. The “un” in the figures is the abbreviation for “unclassified”, and the MM is *Methylobacterium-Methylorubrum*. Several bacterial functions classified as N circles related to the AAL and AAS are shown in Table S1-2. Several fungal guilds classified as plant pathogen-related guilds in the AAL and AAS are shown in Tables S3-4. Red and blue represent negative and positive Spearman’s coefficients, respectively. * *P* < 0.05, ** *P* < 0.01, *** *P* < 0.001. Different lowercase letters in the heatmap represent significant differences in the relative abundance of the same genus between AAS and AAL (*P* < 0.05).

We obtained 192 cultivable fungal isolates from 40 dead seedlings, with an average of 4.825 isolates per dead seedling (Fig. 4a). Based on the ITS genes of the representative strains (Table S5), they were divided into 7 families. The dominant family was Didymellaceae (relative abundance = 66.15%), and the numerically dominant genera were *Allophoma* (50.52%), *Alternaria* (26.04%) and *Epicoccum* (5.73%) (Fig. 4b, Table S5). The seedling-killing effects of these strains on *A. adenophora* exhibited a significant phylogenetic signal (Pagel’s λ = 0.82, *P* = 0.0002). Overall, numerically dominant *Allophoma* (Didymellaceae) and *Alternaria* (Pleosporaceae) had high seedling mortality (54% – 100%) (Fig. 4c, Fig. S3).

**Figure 4.**
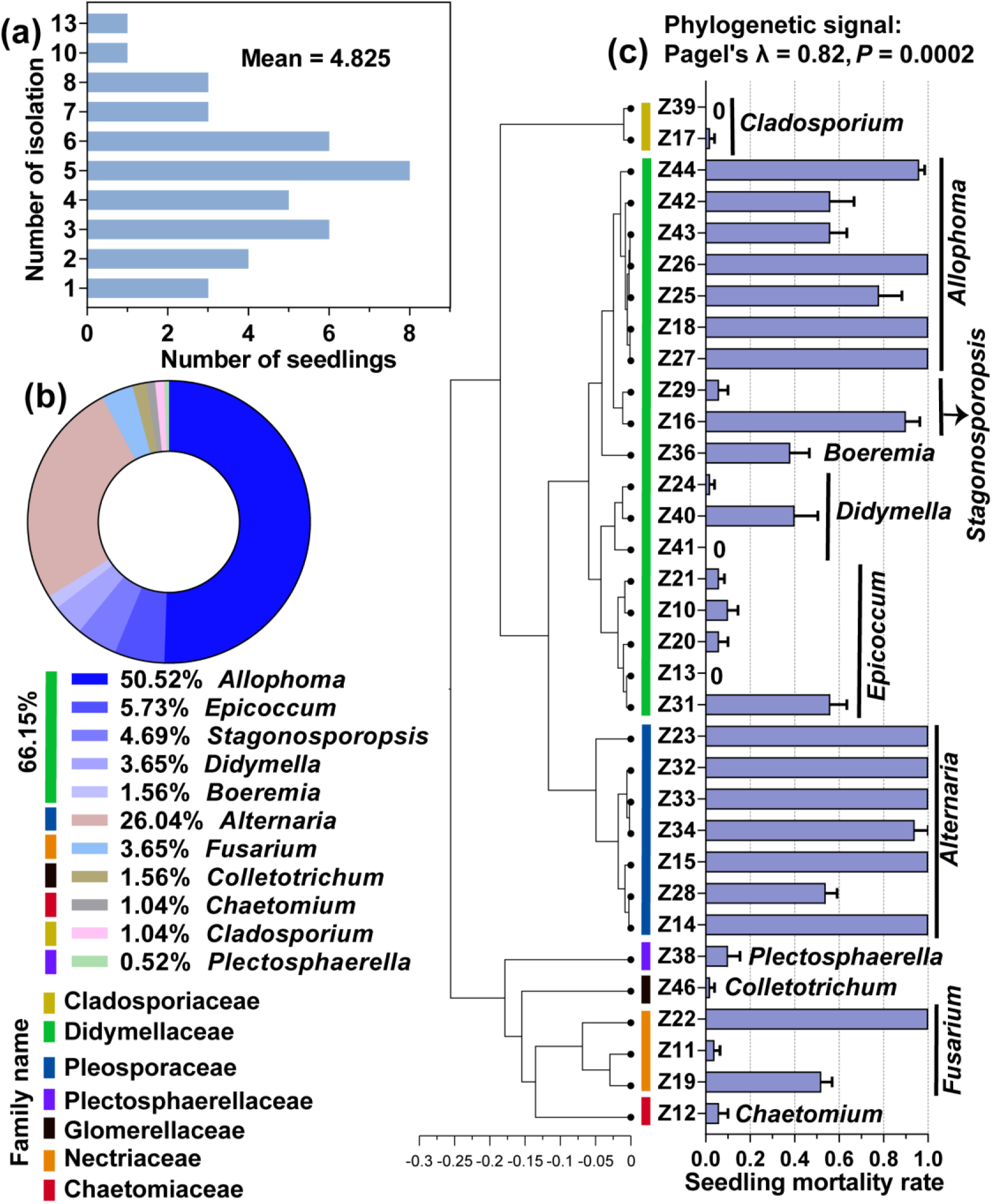
Cultivable fungi associated with dead *A. adenophora* seedlings and their seedling-killing effects on *A. adenophora*. Isolation frequency from one dead seedling (a) and cultivable fungal community composition at the genus level (b). The seedling-killing effects of 33 fungal strains on *A. adenophora* and their phylogenetic signals (c).

### Enrichment of microbial community and function by *A. adenophora* seedlings under different treatments

NMDS and PERMANOVA revealed that all four factors significantly affected the bacterial community and functional assembly of the seedlings, and the greatest effects were inoculation time and compartment (all *P* < 0.05, R^2^: 0.102 – 0.138), followed by inoculation source and nutrition (all *P* < 0.05, R^2^: 0.024 – 0.082, Fig. 5a,b,e,f). Additionally, compartment, inoculation source and time significantly affected the fungal community and functional assembly (all *P* < 0.05, R^2^: 0.054 – 0.102), but nutrition affected only the fungal community (*P* = 0.001, R^2^ = 0.031) (Fig. 5c,d,g,h). Further analysis for each inoculation time treatment showed that compartment and inoculation source mainly affected the microbial community and functional assembly and explained a greater proportion of the variation in bacteria than in fungi and in function than in the community. The nutrient level mainly affected the bacterial community at certain inoculation times (*P* < 0.05 for G0, G21 and G28, Fig. 5i-l, Fig. S4, Table S6-7).

**Figure 5.**
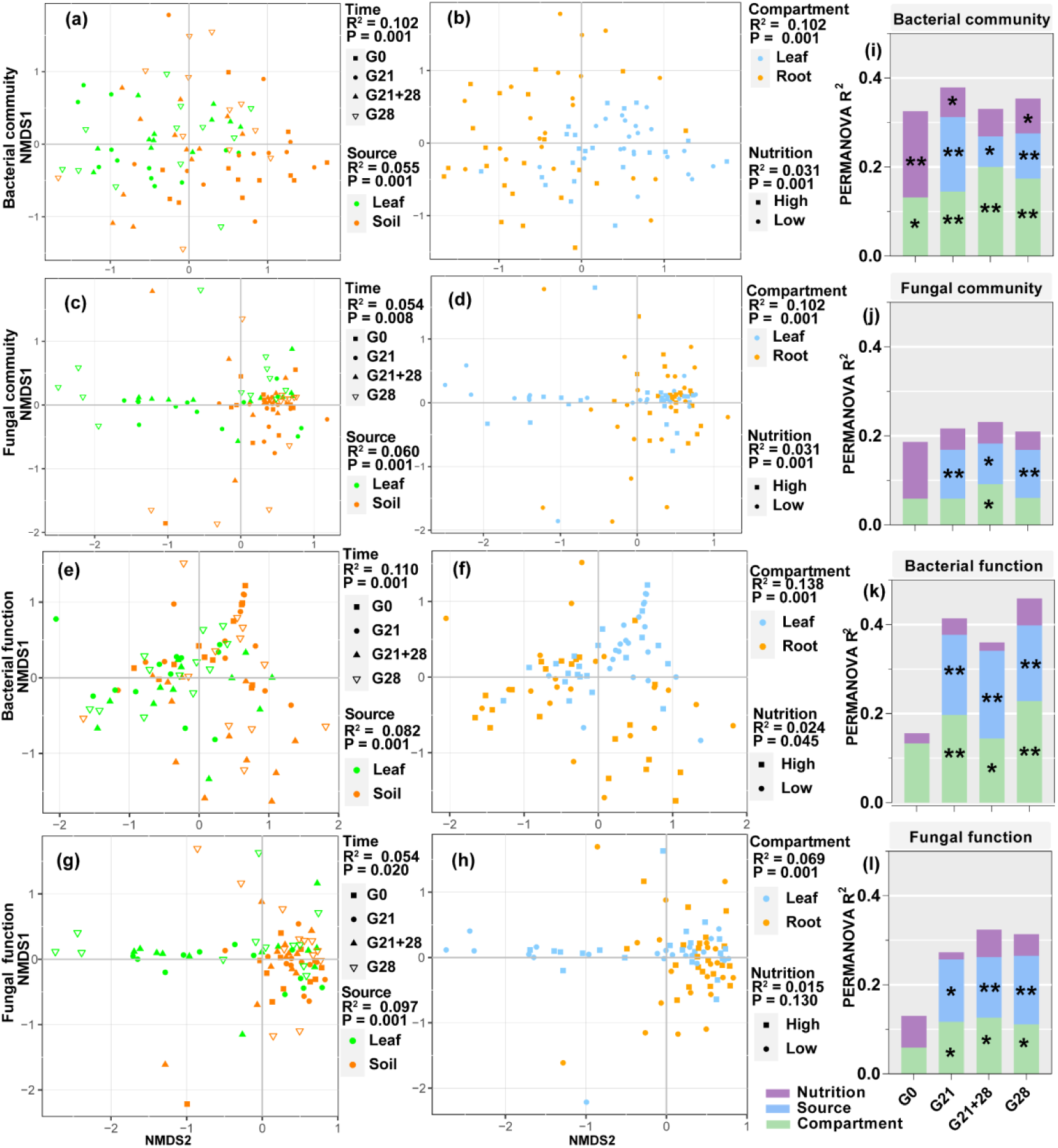
Enrichment of bacterial and fungal communities and functions by *A. adenophora* seedlings under different treatments. Nonmetric multidimensional scaling (NMDS) ordinations of Bray–Curtis dissimilarity matrices with permutational analysis of variance (PERMANOVA) of bacterial and fungal communities and function (a-h). Contribution of plant compartment, inoculation source and nutrient level to the variation in bacterial and fungal communities and function at each inoculation time based on PERMANOVA (i-l). * *P* < 0.05, ** *P* < 0.01, *** *P* < 0.001.

### Correlations of the enriched microbial community and function with *A. adenophora* seedling growth

We further analysed the correlation of microbial abundance and putative functions enriched by seedlings with the microbial effect on seedling growth (RI). We identified 47 root endophytic genera that were significantly correlated with *A. adenophora* growth. Among these genera, seven negative genera were less abundant in seedlings treated by leaf inoculation than in those treated by soil inoculation but were similar in abundance in seedlings treated by different inoculation time, and in seedlings grown at different nutrient levels. In contrast, forty positive genera, e.g., the fungi *Duganella and Mortierella* and the bacteria *Massilia, Pseudomonas, and Sphingomonas*, were more abundant in seedlings treated by leaf inoculation than by soil inoculation but less abundant in seedlings inoculated at G0 than at the other three inoculation times (Fig. 6a). Eighteen leaf endophytic genera with significant correlations with the RI were identified, of which three negative genera, namely, the bacteria *Tardiphaga* and *Brevundimonas* and the fungus *Microsphaera*, were more abundant in seedlings inoculated at G0 than at the other three inoculation time treatments and were slightly more abundant in the seedlings inoculated with soil than in those inoculated with leaf; in contrast, fifteen positive genera, e.g., the fungi *Hypocrea* and Pleosporales_unclassified, were more abundant in the seedlings inoculated with leaf than in those inoculated with soil and were more abundant in the seedlings inoculated at G21 than in the seedlings inoculated at the other three inoculation time treatments (Fig. 6b).

**Figure 6.**
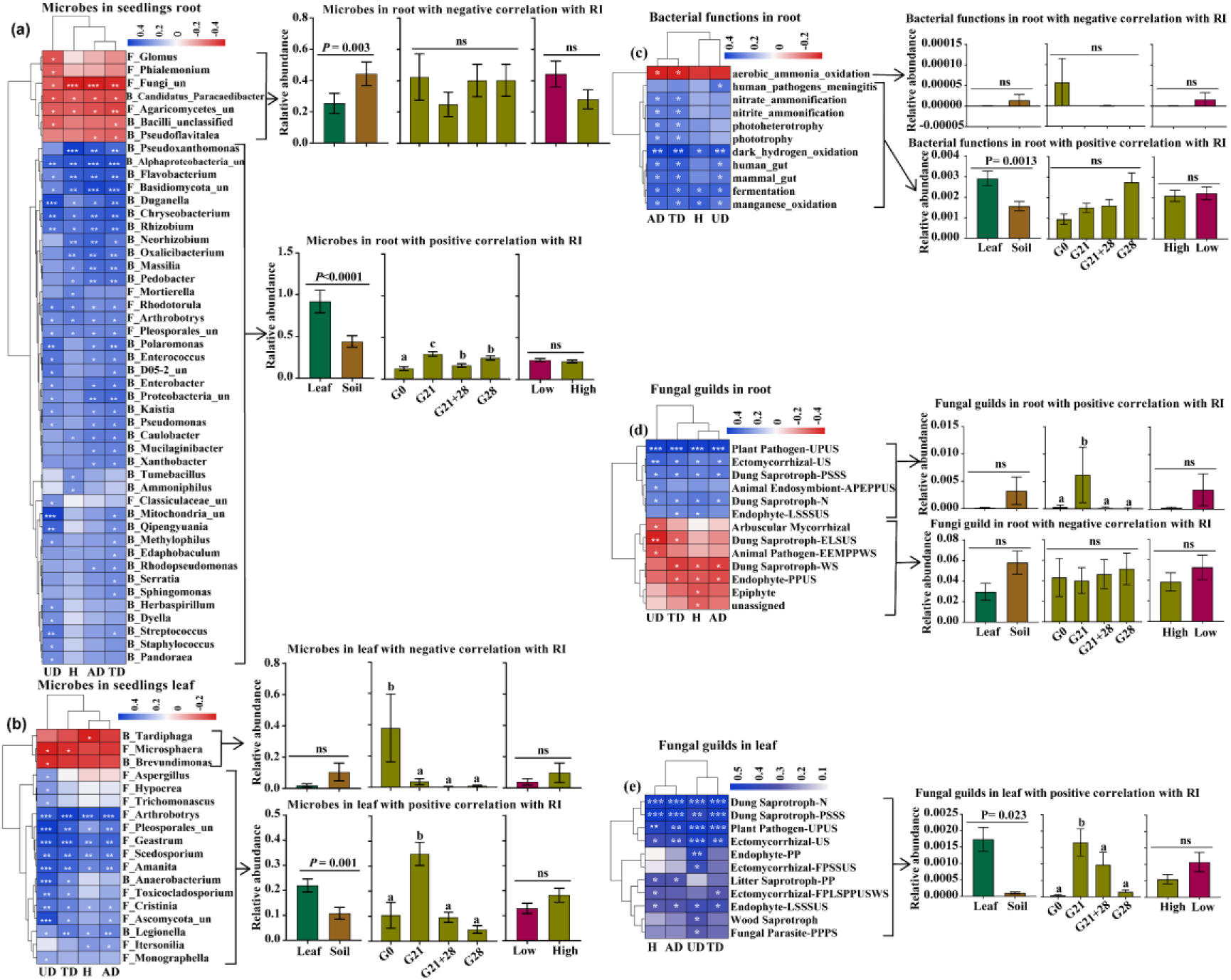
Correlations of the enriched microbial community and function with *A. adenophora* seedling growth. These genera accounted for more than 1% of the total relative abundance in seedling roots or leaves. (a) Correlations between 47 out of 214 genera enriched in roots with RI (left) and the relative abundances of genera under different treatments, i.e., different inoculation sources and time and nutrient levels (right). (b) Correlations between 18 out of 184 genera enriched in leaves with RI (left) and the relative abundances of genera under different treatments (right). (c) Correlations between putative bacterial functions enriched in roots with RIs (left) and the relative abundances of functions with negative and positive correlations under different treatments (right). (d) Correlations between fungal guilds enriched in roots with RIs (left) and the relative abundances of guilds with negative and positive correlations under different treatments. (e) Correlations between fungal guilds in leaves with RIs (left) and the relative abundances of guilds with positive correlations under different treatments (right). No bacterial functions in the leaves showed a significant correlation with seedling growth. “F_” represents fungal genera, “B_” represents bacterial genera, “un” represents unclassified; H: RI of seedling height; AD: RI of aboveground dry biomass; UD: RI of underground dry biomass; TD: RI of total dry biomass. Red and blue represent negative and positive Spearman’s coefficients, respectively. * *P* < 0.05, ** *P* < 0.01, *** *P* < 0.001. For abbreviations of fungal function guilds, please see Table S8.

We identified several bacterial functions in the roots and fungal function guilds enriched in the roots and leaves of *A. adenophora* seedlings that were significantly correlated with the RI (Fig. 6c-e). Two bacterial functions involved in the N cycle (nitrate ammonification and nitrite ammonification) in roots showed a significant positive correlation with RI (Fig. 6c). The fungal guild Ectomycorrhizal showed a significantly positive correlation. Unexpectedly, the putative plant pathogen guild showed a significant positive correlation with seedling growth, while the arbuscular mycorrhizal guild showed a negative correlation (Fig. 6d, e). The positive bacterial functions in roots and fungal guilds in leaves had greater relative abundances in the seedlings after litter inoculation than those after soil inoculation (Fig. 6c, e); additionally, the positive fungal guilds in roots and leaves had significantly greater abundances in the G21 seedlings than in those at the other three inoculation time treatments. Surprisingly, there was no difference in the relative abundance of microbes or functions involved in seedling growth under different nutrient levels (Fig. 6a-e).

## Discussion

### Leaf litter microbes had more adverse effects on *A. adenophora* seed germination and seedling survival than soil microbes

In support of our first expectation, leaf litter had more adverse effects on seed germination and seedling survival than soil (Fig. 1). Leaf litter has been previously reported to have adverse effects on seedling emergence and population establishment (Abbas et al., 2023; Möhler et al., 2021; Zhang et al., 2022) by reducing light and physical barriers and releasing allelochemicals (Abbas et al., 2023; Gross, 1984; Möhler et al., 2018; Zhang et al., 2019). Our study did not directly test the allelopathic effects of leaf litter. However, leaf litter possibly produces allelochemicals that adversely impact *A. adenophora* seed germination time and seedling survival. We observed that sterile leaf litter inoculation caused longer GTs than sterile soil and the control (nothing inoculated) (Fig. 1a). Interestingly, sterile leaf litter inoculation also caused longer GTs than nonsterile leaf litter inoculation, suggesting that some pathways through which leaf microbes alleviate the adverse effects of leaf allelopathy on GTs are unknown. Moreover, sterile leaf inoculation at G0 caused a 19.7% mortality rate for seedlings growing in petri dishes (Fig. 1c), but no dead seedlings were observed when the plants were not inoculated (Fig. 1a, S1).

Nonetheless, our study highlighted the adverse microbial role of leaf litter in seedling mortality because nonsterile leaves have significantly greater seedling mortality (96.7%) than sterile leaves (19.7%) (Fig. 1c). Indeed, we found that leaf litter harbored more abundant bacterial and fungal genera associated with seedling mortality, as well as a greater proportion of plant pathogens, than soil (Fig. 3). The results implied that the litter harboured many plant pathogens and thus played an essential role in mediating *A. adenophora* population density by killing conspecific seedlings. These findings provide novel insights for understanding plant invasion. Invasive plants are commonly characterized by rapid growth and a high yield of seeds (Baker, 1974; Parsons WT, 1992; Zheng et al., 2009). These traits are beneficial for rapid population establishment and range expansion at newly invaded sites. However, these invasive plant species commonly form high-density monocultures once the population is established, e.g., *A. adenophora*. In such a situation, high-density seedlings may exacerbate intraspecific competition. Self-limiting of the population elicited by leaf microbes may in turn help *A. adenophora* maintain monocultures at established sites. Thus, it is highly interesting to determine whether leaf microbe-mediated self-limitation at an early life stage is common and important in other invasive systems.

### Peripheral microbial sources had more adverse effects on seedling survival and growth when inoculating at the early growth stage than at the later stage

Consistent with our second expectation, inoculation time significantly affected seedling survival and growth; in particular, seedling mortality was greater and seedling growth was poorer when inoculated at G0 than at later growth stages (Fig. 1c, d, Fig. 2b). The plant growth stage could change the impact of plant host-associated microbes, and such an impact is always strongest in early plant growth stages (Bagchi et al., 2014; Jevon et al., 2020). One potential reason is that small seedlings usually allocate most of their resources to survive and grow, while older seedlings have relatively more resources to defend against pathogen infection (Geisen et al., 2021; Schloter & Matyssek, 2009). For example, smaller seedlings were more sensitive to inoculated individual fungi, soil microbiota or litter addition than older seedlings due to fewer defense resources and little chance of recovering from biomass loss (Geisen et al., 2021; Jevon et al., 2020; Zhang et al., 2022). Another possible reason is related to the interaction between seed-borne microbes and peripheral microbial sources in young seedlings. There is evidence that seed–borne endophytes are likely to be beneficial for seedling growth and stress resistance (Herrera et al., 2016; Wang & Zhang, 2023). These seed endophytes might be inhibited or even excluded from young seedlings by external sources of microbes inoculated at G0, when seedlings are highly sensitive to inoculated microbes (Geisen et al., 2021; Jevon et al., 2020). Therefore, determining how peripheral microbial sources interact with seed–borne endophytes in seedlings across ontogeny is highly valuable.

We did not observe an adverse effect of leaf litter microbes on *A. adenophora* growth, as observed previously by Kai Fang et al. (2019), who inoculated *A. adenophora* at G0 (sowing seeds), because all seedlings inoculated with leaf litter at G0 died after transplantation into soils in this study. In contrast, both the microbial community and function were significantly positively correlated with seedling growth and had a greater relative abundance in seedlings inoculated with leaf litter than in those inoculated with soil, while those with a negative correlation showed the opposite trend (Fig. 6). This finding suggested that leaf litter microbes might have a more positive effect on *A. adenophora* growth than soil microbes if inoculated during the latter growth stage, such as at G21 and G21+28 (Fig. 2b). Interestingly, we found that N circle-related bacterial functions in seedling roots were positively correlated with seedling growth (Fig. 6c). Similarly, Zhao et al. (2019) showed that *A. adenophora* invasion increased N circle-related bacterial functional genes in soil and subsequently directly promoted plant growth and invasion. K. Fang et al. (2019) also reported that several root endophytic nitrogen–fixing bacteria of *A. adenophora* could significantly promote its growth. Interestingly, the abundance of these N circle–related bacterial functions was greater in seedling roots inoculated with leaf litter than in those inoculated with soil (Fig. 6c), which suggests a possible way in which some N circle–related bacteria associated with leaf litter may migrate from leaves into roots after leaf litter inoculation.

### Nutrient levels did not affect seedling mortality and microbe-mediated *A. adenophora* growth

Nutrient addition can promote severe invasion, as invasive plants often have greater nutrient availability than do native plants. (Shan et al., 2023; Slate et al., 2022). However, it is unclear whether such an advantage is involved in a change in the microbial community-driven host growth effect. We also found that *A. adenophora* grew larger and more rapidly in the high-nutrient treatment than in the low-nutrient treatment (Fig. S6); moreover, the nutrient level significantly changed the microbial community and bacterial function (Fig. 5). Nonetheless, in contrast to our third expectation, seedling mortality was not affected by seedling growth under different nutrient levels, and nutrient levels negligently affected overall microbe-mediated *A. adenophora* growth and the relative abundance of microbes and functions correlated with seedling growth (Figs. 1, 2, 6). Previously, Ai et al. (2018) reported that nutrient additions cause crops to enrich some bacteria and fungi from soil and increase yield; however, there is no evidence that the increased yield effect is due to enriched microbial communities in this study. Our data indicated that the invasion advantage driven by high nutrient availability may be driven primarily by plant physiological traits, such as rapid nutrient absorption and growth strategies, rather than by enriched microbes. Alternatively, it is possible that our delayed harvest of seedlings under low nutrient levels may cover the distinct microbial role in seedling growth between the two nutrient levels (see Methods).

However, there was an interaction effect between nutrient level and inoculation time on seedling growth (Fig. 2). For example, a high nutrient level resulted in a more significant positive microbial effect on seedling growth than a low nutrient level when inoculated at G21, regardless of leaf litter or soil inoculation. It is unclear whether, during the first 21 days before inoculation, more beneficial seed endophytes are enriched to produce a more positive effect on seedling growth under high-nutrition conditions than under low-nutrition conditions, as seed endophytes can facilitate nutrient acquisition and subsequently promote plant growth (Khalaf & Raizada, 2016; Sanchez-Lopez et al., 2018; Shao et al., 2021).

### The same microbial genera had distinct effects on *A. adenophora* seedling survival versus growth

Correlation analysis of the microbial community and function with seedling survival and growth revealed that several genera showed distinct correlations with seedling survival and growth. For example, the bacterial genera *Pseudomonas*, *Sphingomonas* and *Massilia* are positively correlated with seedling mortality and subsequent seedling growth (Figs. 3,6). Many strains belonging to these genera have been reported to promote the growth of many plant species (Jimenez et al., 2020; Luo et al., 2019; Qiao et al., 2019), including *A. adenophora* (Chen et al., 2019; K. Fang et al., 2019), because they are commonly involved in N_2_ fixation (Albino et al., 2006). However, some microbes, e.g., the fungi *Mortierella* and *Hypocrea*, are negatively correlated with early seedling mortality but positively correlated with later seedling growth (Figs. 3, 6). These fungi, as plant growth-promoting fungi (PGPFs), have been widely reported (Contreras-Cornejo et al., 2009; Ozimek & Hanaka, 2021; Wani et al., 2017). These findings suggested that microbial interactions are highly complicated during the early life stage of *A. adeonophora*. On the one hand, there may be sequential effects for some plant growth-promoting microbial groups. For example, the bacteria *Massilia, Pseudomonas* and *Sphingomonas* may negatively affect seedling growth and even kill seedlings if the arrival time is too early after germination. On the other hand, such distinct effects of these bacterial groups on seedling survival versus growth may result from different species from the same genus or even from genetically distinct strains from one species. Interestingly, we found that most *Pseudomonas* and *Sphingomonas* ASVs enriched in the seedlings (>80%) were not associated with the inoculum source (Fig. S8). This suggested that most of the bacterial ASVs positively correlated with growth might be from seeds rather than from the inocula. It is necessary to isolate these enriched microbes to test their interactions with the early life stage of *A. adeonophora*.

Surprisingly, related plant pathogen guilds showed a positive correlation with *A. adenophora* seedling growth (Fig. 6). Because these putative plant pathogens were classified as plant pathogens based on the current database, it remains to determine whether such putative plant pathogens for most native plant species are not detrimental to invader *A. adenophora* growth. Indeed, plant pathogens often can switch from a beneficial endophyte to a pathogen or vice versa depending on different host plant species (Delaye et al., 2013; Tian et al., 2020).

### Seedling-killing microbes were those associated with leaf litter

We found that most seedling-killing microbes isolated from dead seedlings were previously reported to be leaf spot pathogens. For example, *Alternaria* (Pleosporaceae) and several genera belonging to the family Didymellaceae, such as *Allophoma*, *Stagonosporopsis*, *Didymella*, *Boeremia*, and *Epicoccum*, caused high seedling mortality (Fig. 4). *Alternaria* sp. is often pathogenic to a large variety of plants, such as those causing stem cancer, leaf blight or leaf spot (Leiminger et al., 2015; Thomma, 2003; Vergnes et al., 2006), and members of *Allophoma* have also been reported to cause dieback (Babaahmadi et al., 2018) and leaf spot (Garibaldi et al., 2012). All these fungi are leaf spot pathogens of *A. adenophora* and its neighboring native plant (Chen et al., 2022; Kai Fang et al., 2021).

In particular, the numerically dominant *Allophoma* strains obtained in this study had the same ITS genes as the leaf endophyte and leaf spot pathogen *Allophoma* associated with *A. adenophora* (Chen et al., 2022; Kai Fang et al., 2021; Yang et al., 2023). Interestingly, a previous report revealed that the dominant genera in healthy seedlings inoculated with leaf litter were *Didymella* and *Alternaria* (Kai Fang et al., 2019). We did not isolate fungi from healthy seedlings to determine whether the live seedlings indeed lacked or accumulated a lower abundance of the seedling-killing strains than did the dead seedlings in this study. We could assume that these fungal genera likely exist in *A. adenophora* mature individual experiencing a lifestyle switch from endophytic to pathogenic and play an essential role in limiting the population density of *A. adenophora* monocultures by killing seedlings only at very early stages. Thus, it is worth exploring the dynamic abundance of these strains and host resistance variation during *A. adenophora* seedling development.

### Implications for developing biocontrol agents for *A. adenophora* invasion

Our data also have implications for the development of biocontrol agents for *A. adenophora* invasion. Currently, several leaf spot fungi, such as the leaf spot fungus *Phaeoramularia* sp., which is released against *A. adenophora* (Kluge, 1991); the white smut fungus *Entyloma ageratinae* against *A. riparia* (Barton et al., 2007); the rust fungus *Uromycladium tepperianum* against the weed *Acacia saligna* (Wood, 2012); and the rust fungus *Puccinia spegazzinii* against *Mikania micrantha* (Day & Riding, 2019), have been used as biological agents for the control of plant invasion. These agents mainly control weeds by damaging the leaves, stems, and petioles and reducing growth rates, flowering, percentage cover and population density. In this study, the strains associated with leaf litter, such as *Allophoma* sp. and *Alternaria* sp., caused high seedling mortality and thus could control *A. adenophora* invasion at the seedling establishment stage. On the other hand, we found that an external source of microbes had a greater adverse effect on seedling survival and growth when inoculated at G0 than at the later growth stage. Therefore, prevention and control measures by microbial agents taken at the early seedling stage of invasive plants may be more effective than at the mature stage.

## Materials and Methods

### Sample collection and preparation

All seeds, rhizosphere soil (AAS) and leaf litter (AAL) of *A. adenophora* were collected from Xishan Forest Park, Kunming city, Yunnan Province (25°55′34″N; 102°38′30″E, 1890 m), on 9^th^ April 2022. We collected dead leaves (litters) from the stems as inoculated leaves to avoid contamination by soil microbes; moreover, dead leaves could better represent litter in natural systems than fresh leaves. All leaf litter and soil samples were collected from five *A. adenophora* populations ∼200 m away from each other and treated as independent biological replicates. These *A. adenophora* plants had been grown in monoculture for more than 10 years; thus, their rhizosphere soils and leaf litters were used in our feedback experiment rather than via a typical two-phase approach (the first conditioning phase and the second testing phase) (Brinkman et al., 2010). The collected soil and litter samples were naturally dried in a clean room and weighed. The soil was ground to a 2 mm sieve before weighing. For convenience in the inoculation application, we prepared these samples in leaf litter bags (each containing 2 g of leaf litter) (Zaret et al., 2021) and soil bags (each containing 5 g of soil) as well as 0.1 g of soil or litter (cut into pieces smaller than 2×2 mm) in centrifuge tubes. All sample bags and tubes were divided into nonsterile and sterile groups and stored at 4°C until inoculation. The sterile groups (soils or litter) were sterilized by gamma irradiation (30 kGy, 30 h, Huayuan Nuclear Radiation Technology Co., Ltd., Kunming, China), which can kill all microorganisms because no colonies were formed after 7 days of inoculation on PDA media for gamma-irradiated samples (see Fig. S5); moreover, there was no evidence that this irradiation method changed the chemistry of the samples. The nonsterile groups (soils or litter) were natural samples containing live microorganisms. The natural soil or litter (0.3 g) was weighed into tubes and placed at –80°C until DNA extraction.

### Experimental design

The experimental design is shown in Fig. 7. (1) We inoculated *A. adenophora* with nonsterile rhizosphere soil and leaf litter of *A. adenophora* and sterile groups as control groups to distinguish the effects of aboveground microbes from those of underground microbes. (2) We inoculated the soils or leaf litters of *A. adenophora* at three growth stages (0 d, 21 d, 28 d) to explore the susceptibility of the seedlings to microbial infection and growth effects. We performed inoculation on the day of sowing the seeds on water agar plates (containing 5 g of agar and 500 mL of water) for germination, which we named G0 inoculation; on the 21^st^ day after seedling growth, we named G21; and on the 28^th^ day after seedling growth, we named G28. Leaf inoculation at G28 was performed to simulate natural microbial spread from the leaf litter to the above part of the seedlings by suspending the leaf bag over the transplanted seedlings without direct contact all the time (see Zaret et al. (2021)). This method may result in only microbial species with easy air transmission to infect seedlings. Thus, an additional combination inoculation (named G21+28) was performed on both the 21st (with seedling contact) and 28th days (without seedling contact) to ensure that most leaf microbes had the opportunity to reach the seedlings. Therefore, we had four treatments for inoculation time. (3) We grew all the inoculated seedlings in background soil (made of Pindstrup substrate, pearlite and vermiculite at a volume ratio of 8:1:1, and nutrient content of Pindstrup substrate, see Table S9) by adding the same volume of Hoagland nutrient solution (high nutrient level) or only water (low nutrient level) to the soil. In total, our experimental design included 4 inoculation time (G0, G21, G28, G21+28) × 2 inoculum sources (leaf litter, rhizosphere soil) × 2 microbial treatments (sterile, nonsterile) × 2 nutrient levels (high, low) × 5 replicates = 160 cups. All the cups were randomly placed in the growth chamber and rearranged randomly every week to mitigate potential positional effects. Seedlings were harvested after 8 weeks of growth under high-nutrient conditions because they grew too fast and touched the PTFE cover; however, we harvested those plants grown under low-nutritional conditions after another 4 weeks of growth due to their very small size (see Fig. S6). No seedlings survived at the G0 inoculation of nonsterile leaf litters when harvested. Stem height, dry aboveground biomass and underground biomass were measured at harvest. Fresh seedling leaves and roots (0.3 g) from three seedlings per treatment as three replicates were harvested and surface-sterilized for microbial community detection. The aboveground and underground dry biomasses for seedlings with less than 0.3 g fresh weight were obtained by linear regression (see Fig. S7). Seed germination and inoculation manipulation for each inoculation treatment are detailed in supplementary method 1 (also see Fig. 7).

**Figure 7.**
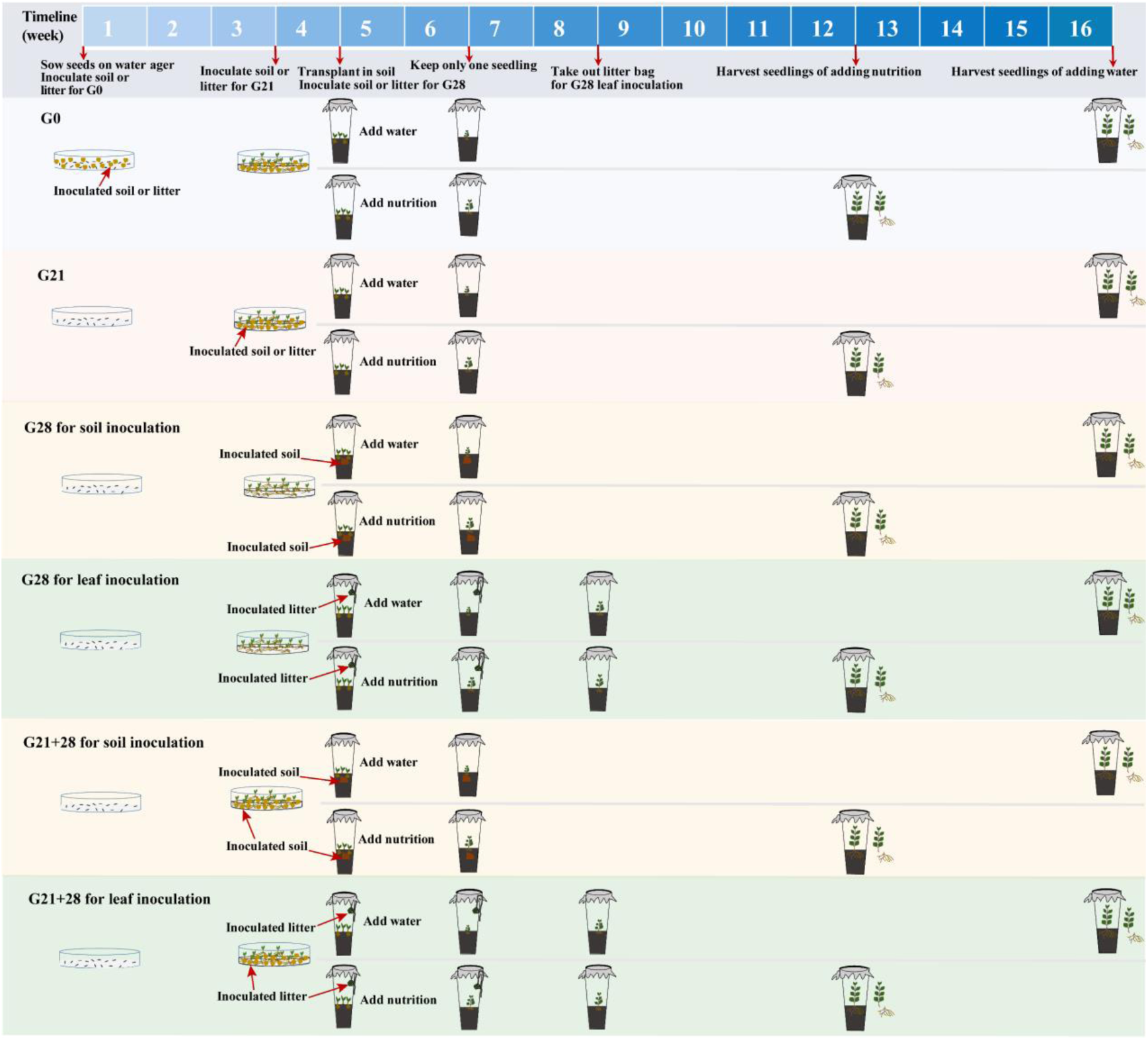
Schematic diagram of soil or litter inoculation at different growth stages. The abbreviations G0, G21, G28, and G21+28 are shown in Fig. 1.

### Molecular sequencing of the microbial community

To link microbial sources (leaf litter and soil) with seed germination, seedling mortality and subsequent seedling biomass, we sequenced the microbial community associated with inoculum samples (natural *A. adenophora* soils (AAS) and leaf litter (AAL)), as well as fresh leaves and roots of *A. adenophora* seedlings grown in the nonsterile treatments. Fresh leaves and roots (0.3 g) of each seedling at harvest were chosen for surface sterilization and then stored at –80°C until total DNA was extracted. For DNA extraction, target‒gene amplification and sequencing steps, see Supplementary Method 2.

### Seedling-killing fungus experiment

No dead seedlings were observed from Petri dishes inoculated with nonsterile soils at G0. Thus, we used 40 dead seedlings obtained from Petri dishes inoculated with nonsterile leaf litter at G0 to isolate fungi. Each dead seedling was cut into 1 mm × 1 mm pieces, and three tissues were placed on each PDA Petri dish and incubated at ambient temperature (20–25°C) for 6–8 days or until mycelia grew. Hyphal tip cultures were subsequently transferred onto new PDA plates and incubated until pure colonies appeared. The DNA of these purified fungal strains was extracted and identified by sequencing the ITS region (for details, see Supplementary Method 2).

To test the seedling-killing effects of these fungal strains on *A. adenophora*, sixteen surface-sterilized *A. adenophora* seeds were sown in a water agar plate in a Petri dish. Ten similar-sized seedlings in one Petri dish 21 days after sowing were selected for fungal inoculation. Five Petri dishes were used as five replicates for each strain. Fungi were grown on PDA for 7 days in an incubator at 25°C, after which 3 mm diameter agar discs with fungal mycelia were inoculated into seedlings by touching the leaves or stems (see Fig. S3). Seedlings were regarded as dead when the leaf and stem became brown and rotten. We recorded the number of dead seedlings after 14 days of inoculation with agar discs and then calculated the mortality rate (MR = the number of dead seedlings/10).

### Statistical analysis

It is unreasonable to directly compare seedling biomass among treatments because of different harvest time under high or low soil nutrition conditions (see methods description above), the response index (RI) was calculated to evaluate the feedback intensity (or direction) of microbes in the inocula soil or leaf on seedling growth: RI = (variable_nonsterile_ – variable_sterile_)/variable_sterile_) (Bruce Williamson & Richardson, 1988), a one‒sample T test was used to determine the significance between the RI value and zero, where RI > 0 and < 0 represent microbes that promote or inhibit seedling growth, respectively. Because the mortality rates of some sterile groups were zero and their RIs were impossible to calculate, we had to directly compare the seedling mortality caused by nonsterile with by sterile samples and perform the analysis of correlation between the mortality rate and microbial composition. Generalized linear models (GLMs) with Gaussian error distributions (identity link) generated by the “lme4” package were used to identify the effects of inoculation source, time, nutrient level treatments and their interaction on the RIs of plant growth. The R^2^ values of the models were obtained by the “piecewiseSEM” package, and P values were estimated using the ANOVA function via chi-squared (χ2) tests in GLMs. The nonparametric Mann–Whitney U test was used to perform all two-group comparisons, and the Kruskal–Wallis test was performed to compare the differences in mortality rate, RI or microbial relative abundances among the four inoculation time treatments.

Nonparametric Mann–Whitney U tests, Kruskal–Wallis tests and one‒sample T tests were performed using SPSS v. 22.0 (SPSS, Inc., Chicago, IL, USA). Nonmetric multidimensional scaling (NMDS) analysis was used to visualize the similarities in bacterial and fungal composition and function among the treatments. Permutational analysis of variance (PERMANOVA) was performed with the ADONIS function in the R (v. 4.2.0) package “vegan” to test the differences in the bacterial and fungal communities and functions among the treatment groups. Bacterial functional profiles were predicted using functional annotation of prokaryotic taxa (FAPROTAX) (Louca et al., 2016). Fungal functional guilds were inferred using the program FUNGuild, and guild assignments with confidence rankings of “Highly probable” and “Probable” were retained (Nguyen et al., 2016). The core microbial taxa were primarily selected from the ASVs that appeared (100% prevalence) among all the samples. Spearman’s correlation analysis was used to link microbial communities in inoculation sources (leaf litter and soils) with seed germination and seedling mortality in Petri dishes of the nonsterile G0 treatment, as well as to link the RI of seedling growth with microbial communities or functions in seedling leaves or roots based on all treatments. A correlation was considered significant when *P* < 0.05. Heatmap plotting was performed in R 4.2.0 with the “pheatmap” package. To examine the phylogenetic signal of the seedling-killing of fungal strains on *A. adenophora*, we calculated Pagel’s λ with the R package “phytools”, which measures the distribution of a trait across a phylogeny. A Pagel’s λ closer to 1 indicated a stronger phylogenetic signal (Pagel, 1999). The remaining figures were visualized in GraphPad Prism v7.0 (GraphPad Software, Inc., San Diego, CA, USA).

## Data availability

The raw bacterial and fungal sequence data are archived in GenBank under accession numbers PRJNA1008375 and PRJNA1008403, respectively. Fungal ITS sequences could be obtained from GenBank under accession numbers OR473386-OR473418.

## Acknowledgements

We thank Xiao-Han Jin, Yu Li, Jin-Peng Li, and Lu Cheng at Yunnan University for their help with sample collection and Jing-Yao Zhang and Ke-Yu Zhou at Yunnan University for their help with the experiments.

## Author contributions

Zhao-ying Zeng: conceptualization, formal analysis, investigation, methodology, visualization, writing – original draft, writing – review & editing. Jun-rong Huang, Zi-qing Liu, Ai-ling Yang, Yu-xuan Li and Yong-lan Wang: investigation. Han-bo Zhang: conceptualization, writing – review & editing, supervision, funding acquisition, project administration.

## References

Abbas, A. M., Alomran, M. M., Alharbi, N. K., & Novak, S. J. (2023). Suppression of Seedling Survival and Recruitment of the Invasive Tree Prosopis juliflora in Saudi Arabia through Its Own Leaf Litter: Greenhouse and Field Assessments. Plants (Basel), 12(4). 10.3390/plants12040959

Ai, C., Zhang, S. Q., Zhang, X., Guo, D. D., Zhou, W., & Huang, S. M. (2018). Distinct responses of soil bacterial and fungal communities to changes in fertilization regime and crop rotation. Geoderma, 319, 156–166. 10.1016/j.geoderma.2018.01.010

Albino, U., Saridakis, D. P., Ferreira, M. C., Hungria, M., Vinuesa, P., & Andrade, G. (2006). High diversity of diazotrophic bacteria associated with the carnivorous plant Drosera villosa var. villosa growing in oligotrophic habitats in Brazil. Plant and Soil, 287(1-2), 199–207. 10.1007/s11104-006-9066-7

Babaahmadi, G., Mehrabi-Koushki, M., & Hayati, J. (2018). Allophoma hayatii sp nov., an undescribed pathogenic fungus causing dieback of Lantana camara in Iran. Mycological Progress, 17(3), 365–379. 10.1007/s11557-017-1360-7

Bagchi, R., Gallery, R. E., Gripenberg, S., Gurr, S. J., Narayan, L., Addis, C. E., Freckleton, R. P., & Lewis, O. T. (2014). Pathogens and insect herbivores drive rainforest plant diversity and composition. Nature, 506(7486), 85-+. 10.1038/nature12911

Baker, H. G. (1974). The Evolution of Weeds. Annual Review of Ecology and Systematics, 5(1), 1–24. 10.1146/annurev.es.05.110174.000245

Barton, J., Fowler, S. V., Gianotti, A. F., Winks, C. J., de Beurs, M., Arnold, G. C., & Forrester, G. (2007). Successful biological control of mist flower (Ageratina riparia) in New Zealand: Agent establishment, impact and benefits to the native flora. Biological Control, 40(3), 370–385. 10.1016/j.biocontrol.2006.09.010

Bauer, J. T., Blumenthal, N., Miller, A. J., Ferguson, J. K., & Reynolds, H. L. (2017). Effects of between-site variation in soil microbial communities and plant-soil feedbacks on the productivity and composition of plant communities. Journal of Applied Ecology, 54(4), 1028–1039. 10.1111/1365-2664.12937

Bever, J. D., Westover, K. M., & Antonovics, J. (1997). Incorporating the Soil Community into Plant Population Dynamics: The Utility of the Feedback Approach. Journal of Ecology, 85(5), 561–573. 10.2307/2960528

Brinkman, E. P., Van der Putten, W. H., Bakker, E. J., & Verhoeven, K. J. F. (2010). Plant-soil feedback: experimental approaches, statistical analyses and ecological interpretations. Journal of Ecology, 98(5), 1063–1073. 10.1111/j.1365-2745.2010.01695.x

Bruce Williamson, G., & Richardson, D. (1988). Bioassays for allelopathy: Measuring treatment responses with independent controls. Journal of Chemical Ecology, 14(1), 181–187. 10.1007/BF01022540

Bunyoo, C., Roongsattham, P., Khumwan, S., Phonmakham, J., Wonnapinij, P., & Thamchaipenet, A. (2022). Dynamic Alteration of Microbial Communities of Duckweeds from Nature to Nutrient-Deficient Condition. Plants-Basel, 11(21), Article 2915. 10.3390/plants11212915

Callaway, R. M., Montesinos, D., Williams, K., & Maron, J. L. (2013). Native congeners provide biotic resistance to invasive Potentilla through soil biota. Ecology, 94(6), 1223–1229. 10.1890/12-1875.1

Chen, L., Fang, K., Zhou, J., Yang, Z. P., Dong, X. F., Dai, G. H., & Zhang, H. B. (2019). Enrichment of soil rare bacteria in root by an invasive plant Ageratina adenophora. Science of the Total Environment, 683, 202–209. 10.1016/j.scitotenv.2019.05.220

Chen, L., Yang, A.-L., Li, Y.-X., & Zhang, H.-B. (2022). Virulence and Host Range of Fungi Associated With the Invasive Plant Ageratina adenophora [Original Research]. Front Microbiol, 13. 10.3389/fmicb.2022.857796

Contreras-Cornejo, H. A., Macías-Rodríguez, L., Cortés-Penagos, C., & López-Bucio, J. (2009). Trichoderma virens, a Plant Beneficial Fungus, Enhances Biomass Production and Promotes Lateral Root Growth through an Auxin-Dependent Mechanism in Arabidopsis. Plant Physiology, 149(3), 1579–1592. 10.1104/pp.108.130369 %J Plant Physiology

Day, M. D., & Riding, N. (2019). Host specificity of Puccinia spegazzinii (Pucciniales: pucciniaceae), a biological control agent for Mikania micrantha (Asteraceae) in Australia. Biocontrol Science and Technology, 29(1), 19–27. 10.1080/09583157.2018.1520807

Delaye, L., Garcia-Guzman, G., & Heil, M. (2013). Endophytes versus biotrophic and necrotrophic pathogens-are fungal lifestyles evolutionarily stable traits? Fungal Diversity, 60(1), 125–135. 10.1007/s13225-013-0240-y

Demey, A., Staelens, J., Baeten, L., Boeckx, P., Hermy, M., Kattge, J., & Verheyen, K. (2013). Nutrient input from hemiparasitic litter favors plant species with a fast-growth strategy. Plant and Soil, 371(1-2), 53–66. 10.1007/s11104-013-1658-4

Dostál, P. (2021). The temporal development of plant-soil feedback is contingent on competition and nutrient availability contexts. Oecologia, 196(1), 185–194. 10.1007/s00442-021-04919-6

Fang, K., Bao, Z. S. N., Chen, L., Zhou, J., Yang, Z. P., Dong, X. F., & Zhang, H. B. (2019). Growth-promoting characteristics of potential nitrogen-fixing bacteria in the root of an invasive plant Ageratina adenophora. Peerj, 7, Article e7099. 10.7717/peerj.7099

Fang, K., Chen, L., Zhou, J., Yang, Z.-P., Dong, X.-F., & Zhang, H.-B. (2019). Plant-soil-foliage feedbacks on seed germination and seedling growth of the invasive plant Ageratina adenophora. Proceedings of the Royal Society B-Biological Sciences, 286(1917), Article 20191520. 10.1098/rspb.2019.1520

Fang, K., Chen, L. M., & Zhang, H. B. (2021). Evaluation of foliar fungus-mediated interactions with below and aboveground enemies of the invasive plant Ageratina adenophora. ECOLOGY AND EVOLUTION, 11(1), 526–535. 10.1002/ece3.7072

Fang, K., Zhou, J., Chen, L., Li, Y.-X., Yang, A.-L., Dong, X.-F., & Zhang, H.-B. (2021). Virulence and community dynamics of fungal species with vertical and horizontal transmission on a plant with multiple infections. PLOS Pathogens, 17(7), e1009769. 10.1371/journal.ppat.1009769

Flory, S. L., & Clay, K. (2013). Pathogen accumulation and long-term dynamics of plant invasions. Journal of Ecology, 101(3), 607–613. 10.1111/1365-2745.12078

Garibaldi, A., Gilardi, G., Ortu, G., & Gullino, M. L. (2012). First Report of Leaf Spot of Lettuce (Lactuca sativa L.) Caused by Phoma tropica in Italy. Plant Disease, 96(9), 1380–1380. 10.1094/pdis-04-12-0394-pdn

Geisen, S., ten Hooven, F. C., Kostenko, O., Snoek, L. B., & van der Putten, W. H. (2021). Fungal root endophytes influence plants in a species-specific manner that depends on plant’s growth stage. JOURNAL OF ECOLOGY, 109(4), 1618–1632. 10.1111/1365-2745.13584

Gross, K. L. (1984). Effects of seed size and growth form on seedling establishment of 6 monocarpic perennial plants. Journal of Ecology, 72(2), 369–387. 10.2307/2260053

Gu, C., Tu, Y., Liu, L., Wei, B., Zhang, Y., Yu, H., Wang, X., Yangjin, Z., Zhang, B., & Cui, B. (2021). Predicting the potential global distribution of Ageratina adenophora under current and future climate change scenarios. ECOLOGY AND EVOLUTION, 11(17), 12092–12113. 10.1002/ece3.7974

Gustafson, D. J., & Casper, B. B. (2004). Nutrient addition affects AM fungal performance and expression of plant/fungal feedback in three serpentine grasses. PLANT AND SOIL, 259(1-2), 9–17. 10.1023/B:PLSO.0000020936.56786.a4

Herrera, S. D., Grossi, C., Zawoznik, M., & Groppa, M. D. (2016). Wheat seeds harbour bacterial endophytes with potential as plant growth promoters and biocontrol agents of Fusarium graminearum. Microbiol Res, 186, 37–43. 10.1016/j.micres.2016.03.002

Jessen, M.-T., Auge, H., Harpole, W. S., & Eskelinen, A. (2023). Litter accumulation, not light limitation, drives early plant recruitment. Journal of Ecology, 111(6), 1174–1187. 10.1111/1365-2745.14099

Jevon, F. V., Record, S., Grady, J., Lang, A. K., Orwig, D. A., Ayres, M. P., & Matthes, J. H. (2020). Seedling survival declines with increasing conspecific density in a common temperate tree. Ecosphere, 11(11), e03292. 10.1002/ecs2.3292

Jimenez, J. A., Novinscak, A., & Filion, M. (2020). Pseudomonas fluorescens LBUM677 differentially increases plant biomass, total oil content and lipid composition in three oilseed crops. Journal of Applied Microbiology, 128(4), 1119–1127. 10.1111/jam.14536

Julien, M. H., & Griffiths, M. W. (1998). Biological control of weeds: a world catalogue of agents and their target weeds.. ed. 4. C.A.B. International.

Kardol, P., Cornips, N. J., van Kempen, M. M. L., Bakx-Schotman, J. M. T., & van der Putten, W. H. (2007). Microbe-mediated plant-soil feedback causes historical contingency effects in plant community assembly. Ecological Monographs, 77(2), 147–162. 10.1890/06-0502

Khalaf, E. M., & Raizada, M. N. (2016). Taxonomic and functional diversity of cultured seed associated microbes of the cucurbit family. BMC microbiology, 16, Article 131. 10.1186/s12866-016-0743-2

Kluge, R. L. (1991). Biological control of crofton weed, Ageratina adenophora (Asteraceae), in South Africa. Agriculture, Ecosystems & Environment, 37(1), 187–191. 10.1016/0167-8809(91)90146-O

Lamb, E. G. (2008). Direct and indirect control of grassland community structure by litter, resources, and biomass. Ecology, 89(1), 216–225. 10.1890/07-0393.1

Leiminger, J., Bassler, E., Knappe, C., Bahnweg, G., & Hausladen, H. (2015). Quantification of disease progression of Alternaria spp. on potato using real-time PCR. European Journal of Plant Pathology, 141(2), 295–309. 10.1007/s10658-014-0542-2

Liu, B., Daryanto, S., Wang, L. X., Li, Y. J., Liu, Q. Q., Zhao, C., & Wang, Z. N. (2017). Excessive Accumulation of Chinese Fir Litter Inhibits Its Own Seedling Emergence and Early Growth-A Greenhouse Perspective. Forests, 8(9), Article 341. 10.3390/f8090341

Liu, Y., Zheng, Y. L., Jahn, L. V., & Burns, J. H. (2023). Invaders responded more positively to soil biota than native or noninvasive introduced species, consistent with enemy escape. Biological Invasions, 25(2), 351–364. 10.1007/s10530-022-02919-y

Louca, S., Parfrey, L. W., & Doebeli, M. (2016). Decoupling function and taxonomy in the global ocean microbiome. SCIENCE, 353(6305), 1272–1277. 10.1126/science.aaf4507

Loydi, A., Eckstein, R. L., Otte, A., & Donath, T. W. (2013). Effects of litter on seedling establishment in natural and semi-natural grasslands: a meta-analysis. Journal of Ecology, 101(2), 454–464. 10.1111/1365-2745.12033

Lu, P., Ma, K., & Sang, W. (2006). Effects of environmental factors on germination and emergence of Crofton weed (Eupatorium adenophorum). Weed Science, 54(3), 452–457. 10.1614/WS 05-174R1.1

Luo, Y., Wang, F., Huang, Y. L., Zhou, M., Gao, J. L., Yan, T. Z., Sheng, H. M., & An, L. Z. (2019). Sphingomonas sp. Cra20 Increases Plant Growth Rate and Alters Rhizosphere Microbial Community Structure of Arabidopsis thaliana Under Drought Stress. Front Microbiol, 10, Article 1221. 10.3389/fmicb.2019.01221

Ma, Z. W., Wang, Y. X., Gu, Y. C., Bowatte, S., Zhou, Q. P., & Hou, F. J. (2020). Effects of Litter Leachate on Plant Community Characteristics of Alpine Grassland in Qinghai Tibetan Plateau. Rangeland Ecology & Management, 73(1), 147–155. 10.1016/j.rama.2019.10.003

Mitchell, C. E., & Power, A. G. (2003). Release of invasive plants from fungal and viral pathogens. Nature, 421(6923), 625–627. 10.1038/nature01317

Möhler, H., Diekötter, T., Bauer, G. M., & Donath, T. W. (2021). Conspecific and heterospecific grass litter effects on seedling emergence and growth in ragwort (Jacobaea vulgaris). PLoS One, 16(2), e0246459. 10.1371/journal.pone.0246459

Möhler, H., Diekötter, T., Herrmann, J. D., & Donath, T. W. (2018). Allelopathic vs. autotoxic potential of a grassland weed—evidence from a seed germination experiment. Plant Ecology & Diversity, 11(4), 539–549. 10.1080/17550874.2018.1541487

Nguyen, N. H., Song, Z. W., Bates, S. T., Branco, S., Tedersoo, L., Menke, J., Schilling, J. S., & Kennedy, P. G. (2016). FUNGuild: An open annotation tool for parsing fungal community datasets by ecological guild. FUNGAL ECOLOGY, 20, 241–248. 10.1016/j.funeco.2015.06.006

Niu, H. B., Liu, W. X., Wan, F. H., & Liu, B. (2007). An invasive aster (Ageratina adenophora) invades and dominates forest understories in China: altered soil microbial communities facilitate the invader and inhibit natives. Plant and Soil, 294(1-2), 73–85. 10.1007/s11104-007-9230-8

Olson, B. E., & Wallander, R. T. (2002). Effects of invasive forb litter on seed germination, seedling growth and survival. Basic and Applied Ecology, 3(4), 309–317. 10.1078/1439-1791-00127

Ozimek, E., & Hanaka, A. (2021). Mortierella Species as the Plant Growth-Promoting Fungi Present in the Agricultural Soils. Agriculture-Basel, 11(1), 7. https://www.mdpi.com/2077-0472/11/1/7

Pagel, M. (1999). Inferring the historical patterns of biological evolution. Nature, 401(6756), 877–884. 10.1038/44766

Parsons WT, C. E. (1992). Noxious weeds of Australia. CSIRO Publishing.

Qiao, C. C., Penton, C. R., Xiong, W., Liu, C., Wang, R. F., Liu, Z. Y., Xu, X., Li, R., & Shen, Q. R. (2019). Reshaping the rhizosphere microbiome by bio-organic amendment to enhance crop yield in a maize-cabbage rotation system. APPLIED SOIL ECOLOGY, 142, 136–146. 10.1016/j.apsoil.2019.04.014

Sanchez-Lopez, A. S., Thijs, S., Beckers, B., Gonzalez-Chavez, M. C., Weyens, N., Carrillo-Gonzalez, R., & Vangronsveld, J. (2018). Community structure and diversity of endophytic bacteria in seeds of three consecutive generations of Crotalaria pumila growing on metal mine residues. Plant and Soil, 422(1-2), 51–66. 10.1007/s11104-017-3176-2

Schloter, M., & Matyssek, R. (2009). Tuning growth versus defence–belowground interactions and plant resource allocation. Plant and Soil, 323(1), 1–5. 10.1007/s11104-009-0070-6

Shan, L. P., Oduor, A. M. O., Huang, W., & Liu, Y. J. (2023). Nutrient enrichment promotes invasion success of alien plants via increased growth and suppression of chemical defenses. Ecological Applications. 10.1002/eap.2791

Shao, J. H., Miao, Y. Z., Liu, K. M., Ren, Y., Xu, Z. H., Zhang, N., Feng, H. C., Shen, Q. R., Zhang, R. F., & Xun, W. B. (2021). Rhizosphere microbiome assembly involves seed-borne bacteria in compensatory phosphate solubilization. SOIL BIOLOGY & BIOCHEMISTRY, 159, Article 108273. 10.1016/j.soilbio.2021.108273

Slate, M. L., Matallana-Mejia, N., Aromin, A., & Callaway, R. M. (2022). Nitrogen addition, but not pulse frequency, shifts competitive interactions in favor of exotic invasive plant species. Biological Invasions, 24(10), 3109–3118. 10.1007/s10530-022-02833-3

Teste, F. P., Kardol, P., Turner, B. L., Wardle, D. A., Zemunik, G., Renton, M., & Laliberte, E. (2017). Plant-soil feedback and the maintenance of diversity in Mediterranean-climate shrublands. SCIENCE, 355(6321), 173-+. 10.1126/science.aai8291

Thomma, B. P. (2003). Alternaria spp.: from general saprophyte to specific parasite. Mol Plant Pathol, 4(4), 225–236. 10.1046/j.1364-3703.2003.00173.x

Tian, B., Xie, J., Fu, Y., Cheng, J., Li, B., Chen, T., Zhao, Y., Gao, Z., Yang, P., Barbetti, M. J., Tyler, B. M., & Jiang, D. (2020). A cosmopolitan fungal pathogen of dicots adopts an endophytic lifestyle on cereal crops and protects them from major fungal diseases. Isme j, 14(12), 3120–3135. 10.1038/s41396-020-00744-6

van der Putten, W. H., Bardgett, R. D., Bever, J. D., Bezemer, T. M., Casper, B. B., Fukami, T., Kardol, P., Klironomos, J. N., Kulmatiski, A., Schweitzer, J. A., Suding, K. N., Van de Voorde, T. F. J., & Wardle, D. A. (2013). Plant-soil feedbacks: the past, the present and future challenges. JOURNAL OF ECOLOGY, 101(2), 265–276. 10.1111/1365-2745.12054

Vergnes, D. M., Renard, M. E., Duveiller, E., & Maraite, H. (2006). Identification of Alternaria spp. on wheat by pathogenicity assays and sequencing. Plant Pathology, 55(4), 485–493. 10.1111/j.1365-3059.2006.01391.x

Wang, Y.-L., & Zhang, H.-B. (2023). Assembly and Function of Seed Endophytes in Response to Environmental Stress. Journal of microbiology and biotechnology, 33(9), 1–11. 10.4014/jmb.2303.03004

Wang, Z. N., Wang, D. Y., Liu, Q. Q., Xing, X. S., Liu, B., Jin, S. F., & Tigabu, M. (2022). Meta-Analysis of Effects of Forest Litter on Seedling Establishment. Forests, 13(5), Article 644. 10.3390/f13050644

Wani, Z. A., Kumar, A., Sultan, P., Bindu, K., Riyaz-Ul-Hassan, S., & Ashraf, N. (2017). Mortierella alpina CS10E4, an oleaginous fungal endophyte of Crocus sativus L. enhances apocarotenoid biosynthesis and stress tolerance in the host plant. Scientific Reports, 7, Article 8598. 10.1038/s41598-017-08974-z

Whitaker, B. K., Bauer, J. T., Bever, J. D., & Clay, K. (2017). Negative plant-phyllosphere feedbacks in native Asteraceae hosts - a novel extension of the plant-soil feedback framework. Ecol Lett, 20(8), 1064–1073. 10.1111/ele.12805

Widdig, M., Heintz-Buschart, A., Schleuss, P. M., Guhr, A., Borer, E. T., Seabloom, E. W., & Spohn, M. (2020). Effects of nitrogen and phosphorus addition on microbial community composition and element cycling in a grassland soil. SOIL BIOLOGY & BIOCHEMISTRY, 151, Article 108041. 10.1016/j.soilbio.2020.108041

Wood, A. R. (2012). Uromycladium tepperianum (a gall-forming rust fungus) causes a sustained epidemic on the weed Acacia saligna in South Africa. Australasian Plant Pathology, 41(3), 255–261. 10.1007/s13313-012-0126-6

Xiong, S., & Nilsson, C. (1999). The effects of plant litter on vegetation: a meta-analysis. Journal of Ecology, 87(6), 984–994. 10.1046/j.1365-2745.1999.00414.x

Xu, C. W., Yang, M. Z., Chen, Y. J., Chen, L. M., Zhang, D. Z., Mei, L., Shi, Y. T., & Zhang, H. B. (2012). Changes in non-symbiotic nitrogen-fixing bacteria inhabiting rhizosphere soils of an invasive plant Ageratina adenophora. APPLIED SOIL ECOLOGY, 54, 32–38. 10.1016/j.apsoil.2011.10.021

Xu, L., Naylor, D., Dong, Z. B., Simmons, T., Pierroz, G., Hixson, K. K., Kim, Y. M., Zink, E. M., Engbrecht, K. M., Wang, Y., Gao, C., DeGraaf, S., Madera, M. A., Sievert, J. A., Hollingsworth, J., Birdseye, D., Scheller, H. V., Hutmacher, R., Dahlberg, J., … Coleman-Derr, D. (2018). Drought delays development of the sorghum root microbiome and enriches for monoderm bacteria. Proceedings of the National Academy of Sciences of the United States of America, 115(18), E4284–E4293. 10.1073/pnas.1717308115

Yang, A.-L., Chen, L., Cheng, L., Li, J.-P., Zeng, Z.-Y., & Zhang, H.-B. (2023). Two Novel Species of Mesophoma gen. nov. from China. Current Microbiology, 80(4), 129. 10.1007/s00284-023-03238-8

Zaret, M. M., Bauer, J. T., Clay, K., & Whitaker, B. K. (2021). Conspecific leaf litter induces negative feedbacks in Asteraceae seedlings. Ecology, 102(12), e03557. 10.1002/ecy.3557

Zhang, A., Wang, D., & Wan, S. Q. (2019). Litter addition decreases plant diversity by suppressing seeding in a semiarid grassland, Northern China. Ecology and Evolution, 9(17), 9907–9915. 10.1002/ece3.5532

Zhang, R., Hu, X., Baskin, J. M., Baskin, C. C., & Wang, Y. (2017). Effects of Litter on Seedling Emergence and Seed Persistence of Three Common Species on the Loess Plateau in Northwestern China. Front Plant Sci, 8, 103. 10.3389/fpls.2017.00103

Zhang, X., Ni, X., Heděnec, P., Yue, K., Wei, X., Yang, J., & Wu, F. (2022). Litter facilitates plant development but restricts seedling establishment during vegetation regeneration. Functional Ecology, 36(12), 3134–3147. 10.1111/1365-2435.14200

Zhang, Z. J., Liu, Y. J., Brunel, C., & van Kleunen, M. (2020). Evidence for Elton’s diversity-invasibility hypothesis from belowground. Ecology, 101(12). 10.1002/ecy.3187

Zhao, M. X., Lu, X. F., Zhao, H. X., Yang, Y. F., Hale, L., Gao, Q., Liu, W. X., Guo, J. Y., Li, Q., Zhou, J. Z., & Wan, F. H. (2019). Ageratina adenophora invasions are associated with microbially mediated differences in biogeochemical cycles. Science of the Total Environment, 677, 47–56. 10.1016/j.scitotenv.2019.04.330

Zheng, Y. L., Feng, Y. L., Liu, W. X., & Liao, Z. Y. (2009). Growth, biomass allocation, morphology, and photosynthesis of invasive Eupatorium adenophorum and its native congeners grown at four irradiances. Plant Ecology, 203(2), 263–271. 10.1007/s11258-008-9544-5

