## supplemental figures S1-S8;supplemental tables 1-9 for "Distinct effects of phyllosphere and rhizosphere microbes on invader *Ageratina adenophora* during its early life stages"

#### 1. Supplementary methods

**Method S1: Detailed seed germination and inoculation manipulation for each inoculation time treatment.**

In May 2022, all the seeds were surface sterilized, germinated and subsequently grown in RXZ-380D growth chambers (Ningbo Southeast Instrument Co., Ltd., Ningbo, China) at a temperature of 25/20°C (day/night), a light intensity of 12 000 lux, a 12 h photoperiod and a humidity of 65%.

For G0, we exposed sixteen surface-sterilized seeds to litter or rhizosphere soil from *A. adenophora* when the seeds were sown on a water ager plate. Soils or leaves (0.1 g) were distributed in a thin layer on a plate in 90 mm Petri dishes. Each treatment was replicated five times, resulting in a total of 20 plates (2 inoculum sources × 2 microbial treatments × 5 replicates). We recorded germinated seeds every day from the first to 14$^{th}$ day and on the 21$^{st}$ day after sowing to calculate the germination time (GT), germination rate (GR), and number of dead seedlings on the 28$^{th}$ day. The GT was calculated by the formula GT= $\Sigma(Gi\times i)/\Sigma Gi$ (i: number of days between seed sowing (day 0) and seed germination; Gi: number of seeds germinated on day i), according to (Zhang et al., 2014). The GR was calculated as the proportion of germinated seeds on the 21st day after sowing relative to the total number of sown seeds. The seedling mortality rate (MR) was calculated as the number of dead seedlings divided by the total number of germinated seedlings.

For G21, sixteen surface-sterilized seeds were germinated for 21 days in a water agar plate without soil or litter, after which the number of germinated seedlings was recorded. Plates with more than ten germinated seedlings were chosen for inoculation of 0.1 g of litter or rhizosphere soil for a total of 20 plates (2 inoculum sources × 2 microbial treatments × 5 replicates). After 7 days of inoculation, the numbers of germinated and dead seedlings were recorded to calculate the MR.

For G28, sixteen surface-sterilized seeds were germinated for 28 days on a water agar plate without soil or litter, and the number of germinated seeds every day from the first to 14th day and on the 21st day after sowing were recorded to calculate the germination time (GT) and germination rate (GR) as the control (nothing inoculated). Litter and rhizosphere soil were inoculated when similar-sized seedlings were transplanted into cups after 28 days of germination without soil or litter. For leaf inoculation, 2 g of litter bags was suspended above the plants inside individual cups using twine when three plants were transplanted into the soil in a cup, referring to the study of Zaret et al. (2021), such litter had no direct contact with the soils or plants. Litter (2 g) was placed into individual mesh bags made from 10×15 cm cheese cloth squares. Litter bags were removed after 4 weeks to avoid the effects of extra allelochemicals. For soil inoculation, 5 g of rhizosphere soil was added to 65 g of sterile background soil when three seedlings were transplanted into the soil in a cup.

Inoculation of G21+28 is a combination of G21 and G28. Briefly, 0.1 g of litter or rhizosphere soil was inoculated in a plate after 21 days of surface-sterilized seed germination without soil or litter for a total of 20 plates (2 inoculum sources × 2 microbial treatments × 5 replicates). Seedlings were transplanted into the soil in the cups after 7 days of inoculation, and the litter bags were suspended above the plants inside individual cups as part of continuous leaf inoculation. Litter bags were removed after 4 weeks to avoid the effects of extra allelochemicals. Rhizosphere soil (5 g) inoculum was added to 65 g of sterile background soil for soil inoculation. The graphics for all inoculation time treatments are shown in Fig. 7.

Seedlings from the four inoculation timepoint treatments were transplanted into 1000 mL sterile polypropylene cups containing 65 g of sterile background soil and 120 mL of sterile tap water after 28 days of growth in petri dishes after sowing. Three similar-sized seedlings (approximately 1 cm high with 4 small leaves) were transplanted into each cup and subsequently thinned to one seedling per cup 2 weeks after transplantation to avoid intraspecific competition. We recorded the number of dead seedlings in each cup before thinning. The cups were sealed with PTFE bacterial filter membranes to prevent airborne microbe infection and minimize cross contamination between treatments. The background soil was sterilized by autoclaving three times for 2 h, with a 1-day rest period between cycles. The same volume of water or Hoagland nutrient solution was added after the seedlings were transplanted into cups if needed until seedling harvesting.

##### Method S2: DNA extraction, target–gene amplification and sequencing

Total DNA of soil and plant tissue was extracted using the cetyltrimethylammonium bromide (CTAB) method (Stewart & Via, 1993). The quality of the extracted DNA was assessed via electrophoresis in a 1.5% agarose gel using an ND-1000 spectrophotometer (NanoDrop Technology, Wilmington, USA). A Qubit dsDNA HS assay kit (Invitrogen, USA) was used to quantify the DNA concentration. We amplified the bacterial 16S rDNA V4 region and fungal ITS2 region with the primer sets 515F/806R and ITS1F/ITS4, respectively. PCR amplification was performed in a 50 μL mixture containing 12.5 μL of 2X Phanta Max master mix (Thermo Scientific), 2.5 μL of forward primer, 2.5 μL of reverse primer, 50 ng of DNA as a template, and 25 μL of sterile ddH2O. The PCR conditions for the bacterial 16S rRNA genes were as follows: initial denaturation at 98°C for 30 s; 35 cycles of 98°C for 10 s, 54°C for 30 s, and 72°C for 45 s; and a final extension at 72°C for 10 min. For the fungal ITS2 region, the PCR conditions consisted of an initial denaturation at 98°C for 30 s, 32 cycles of denaturation at 98°C for 10 s and annealing at 54°C for 30 s, and extension at 72°C for

45 s, and a final extension at 72°C for 10 min. The PCR products were purified with AMPure XT beads (Beckman Coulter Genomics, Danvers, MA, USA) and quantified with a Qubit (Invitrogen, USA). The amplicon pools were prepared for sequencing, and the size and quantity of the amplicon library were assessed on an Agilent 2100 Bioanalyzer (Agilent, USA) and with the Library Quantification Kit for Illumina (Kapa Biosciences, Woburn, MA, USA), respectively. The libraries were sequenced on a NovaSeq 6000 platform at LC-BIO Biotech Ltd. (Hangzhou, China). High-quality sequences were obtained after removal of low-quality sequences (quality score < 20 and sequence length < 100 bp). Chimeric sequences were filtered using Vsearch software (v2.3.4). After dereplication using DADA2, we obtained an amplicon sequence variant (ASV) feature table and feature sequence. The ASV sequences with poor alignment performance and singleton ASVs were discarded. Taxonomic identification of bacteria and fungi was performed against the SILVA (v138) (Quast et al., 2013) and UNITE (v8.0) databases (Nilsson et al., 2018), respectively. Alpha diversity was calculated by QIIME2, in which the same number of sequences were extracted randomly by reducing the number of sequences to the minimum of some samples. All the sequences obtained in this study have been deposited in the National Center for Biotechnology Information (NCBI) GenBank under SRA accession number PRJNA1008375 for the bacterial 16S rRNA genes and PRJNA1008403 for the fungal ITS2 genetic region.

Fungal mycelia DNA was also extracted using the cetyltrimethylammonium bromide (CTAB) method (Stewart & Via, 1993). We amplified the ITS region of the fungal DNA with the primers ITS4 and ITS5. PCR was performed in a Veriti 96 Well Thermal Cycler (Applied Biosystems Inc., Foster City, CA, USA) in a 50 reactions volume composed of 25 μL of 2 × PCR Master Mix, 1 μL of each primer (10 μM), 22 μL of ddH2O and 1 μL of template DNA. The PCR conditions consisted of initial denaturation at 94°C for 1 min; 35 cycles of denaturation at 94°C for 1 min, annealing at 54°C for 1 min, and extension at 72°C for 1 min; and a final extension at 72°C for 10 mins. PCR products were purified, and forward amplicons were sequenced by

Sangon Biotech Co., Ltd. (Shanghai, China).

The obtained sequences were edited using EDITSEQ and SEQMAN software in the DNASTAR package (DnaStarInc., Madison, WI, USA). We aligned sequences in MEGA v. 6.0 using MUSCLE with default parameters (Edgar, 2004; Tamura et al., 2013), followed by manual checking of alignments. Taxonomic identification was performed against the UNITE (v8.0) database, and BLASTN analyses were performed against the GenBank database. The ITS sequences reported in this study have been deposited in the GenBank database (for accession numbers, see Table S5). BEAST v. 1.10.4 was used to construct a Bayesian phylogenetic tree (Drummond et al., 2012). The resulting tree was visualized in FigTree v.1.4.3.

### 2. Supplementary figure

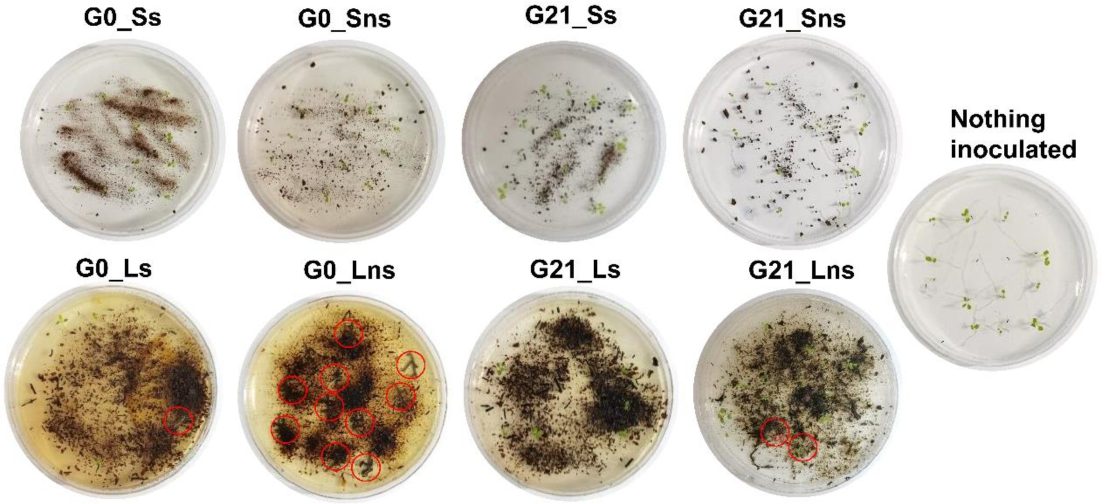


**Figure S1. Germination and dead seedlings of *A. adenophora* inoculated with soil (upper) or litter (bottom) in Petri dishes.** Dead seedlings are circled in red. Ss: sterile soil; Sns: nonsterile soil; Ls: sterile leaf; Lns: nonsterile leaf; G0: inoculated on the day of germination; G21: inoculated on the 21st day after germination. The diameter of the Petri dish is 90 mm.

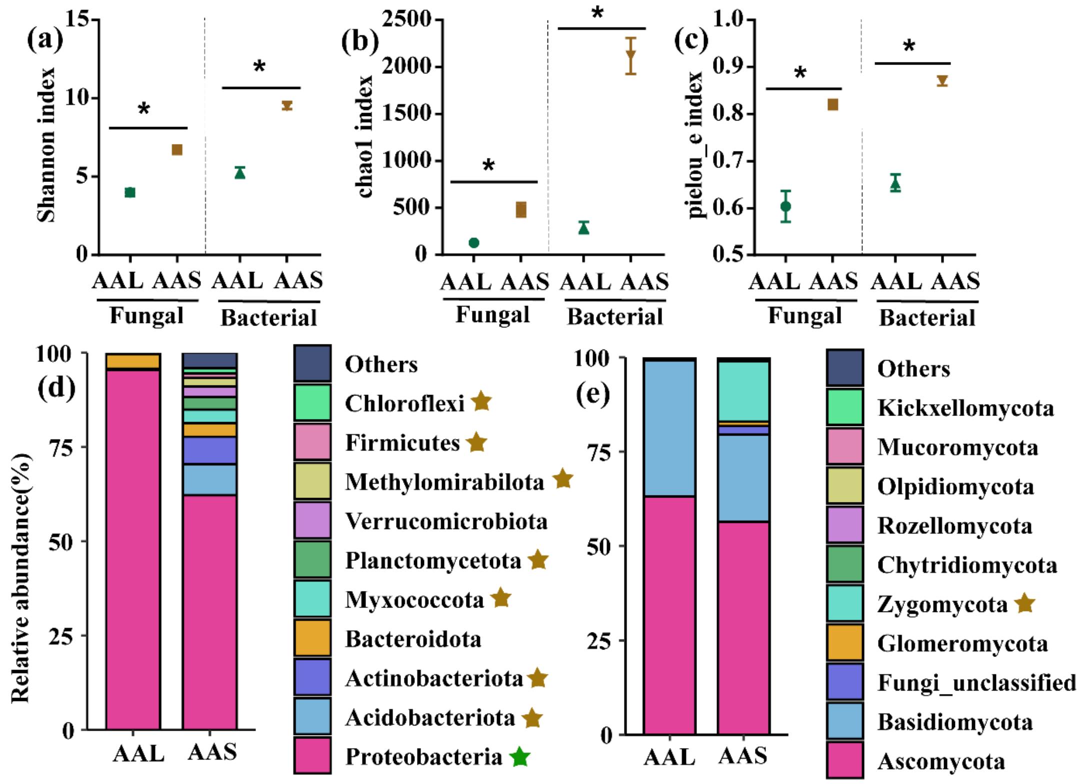


**Figure S2. Microbial diversity and community composition at the phylum level in the inocula AAS and AAL groups.** Fungal and bacterial Shannon, Chao 1 and Pielou_e indices of AAS and AAL (a-c). Relative abundances of bacterial (d) and fungal phyla (e) in the AAS and AAL groups. AAS: *A. adenophora* rhizosphere soil inoculum, AAL: *A. adenophora* litter inoculum. Brown stars represent significantly greater relative abundances in the AAS than in the AAL, and green stars represent significantly greater relative abundances in the AAL than in the AAS, $P < 0.05$ at least.

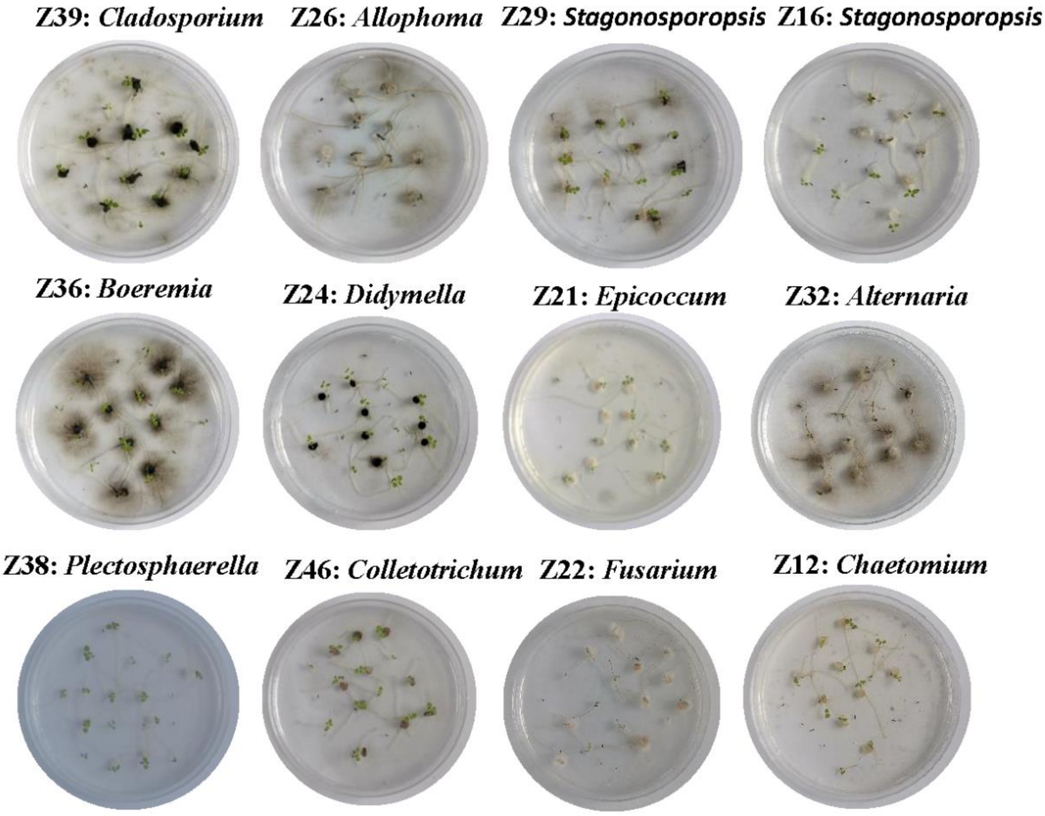


**Figure S3. Seedling-killing effects of several representative strains on *A. adenophora*.** The diameter of the Petri dish is 9 mm.

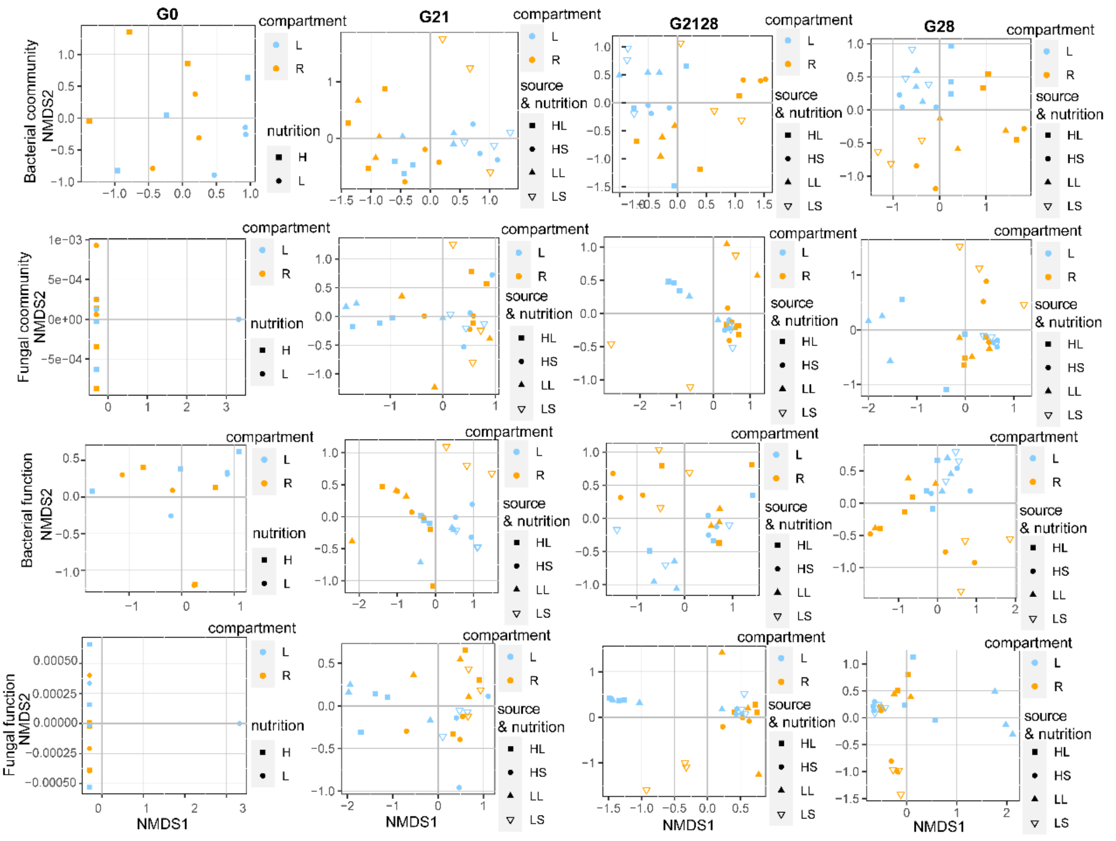


**Figure S4. NMDS results for the bacterial and fungal communities and their functions at each inoculation time treatment.** The different colours represent the different compartments (light blue represents leaves, and orange represents roots). L: leaf; R: root; H: high-nutrition level; L: low-nutrition level; HL: seedling after leaf inoculation under high-nutrition conditions; HS: seedling after soil inoculation under high-nutrition conditions; LL: seedling after leaf inoculation under low-nutrition conditions; LS: seedling after soil inoculation under low-nutrition conditions.

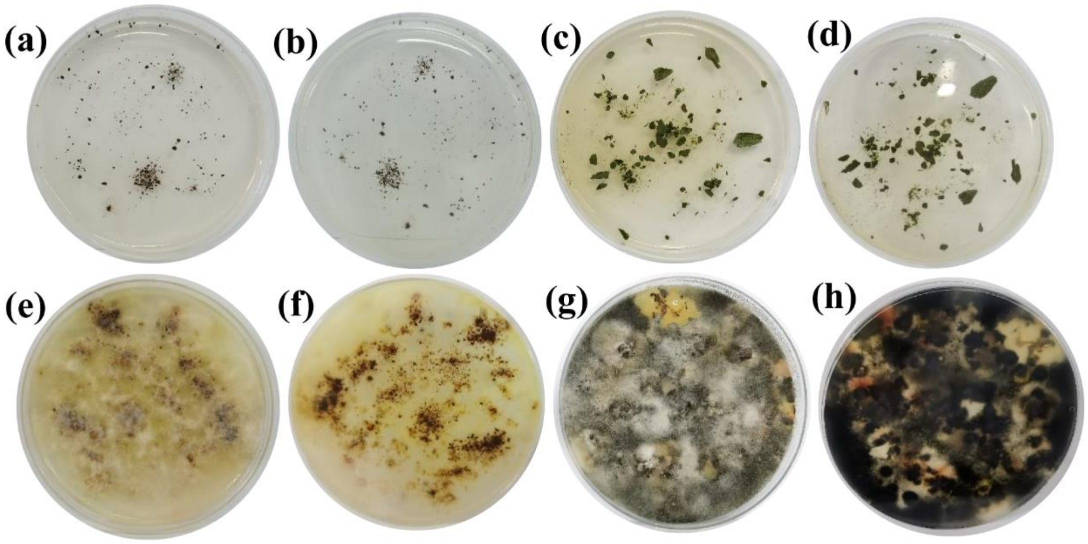


**Figure S5. Gamma irradiation was effective at killing all the microorganisms.** a-d: Front and reverse views of gamma-irradiated soil or leaves inoculated on PDA media on the 7th day. e-h: Front and reverse views of living soil and leaves inoculated on PDA media on the 7th day. The diameter of the Petri dish is 9 mm.

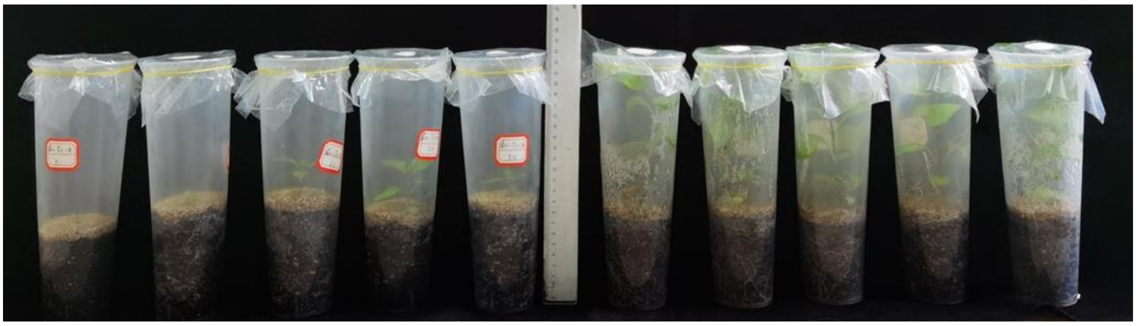

**Figure S6. Seedlings were grown for 8 weeks under low (adding water, left) or high (adding Hoagland nutrient solution, right) nutrient levels.**

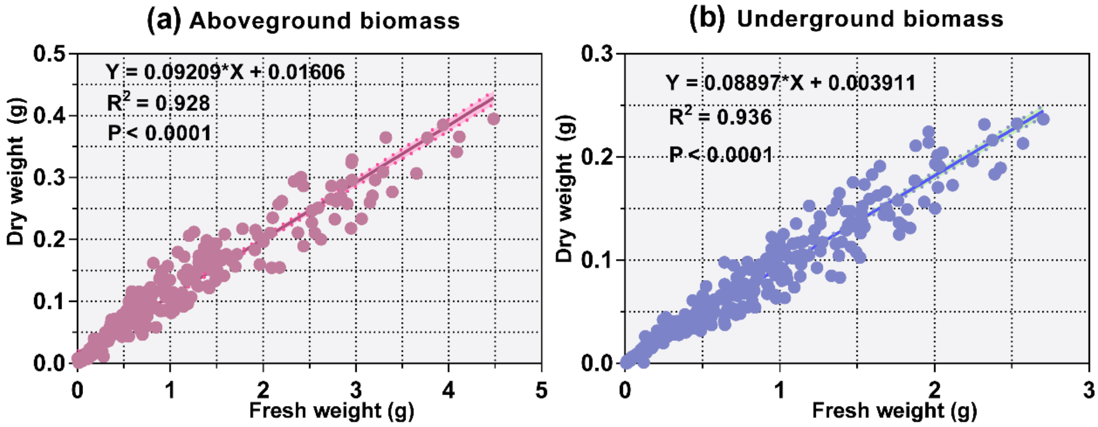


**Figure S7. Linear regression between the fresh and dry weights of aboveground biomass (a) and underground biomass (b).**

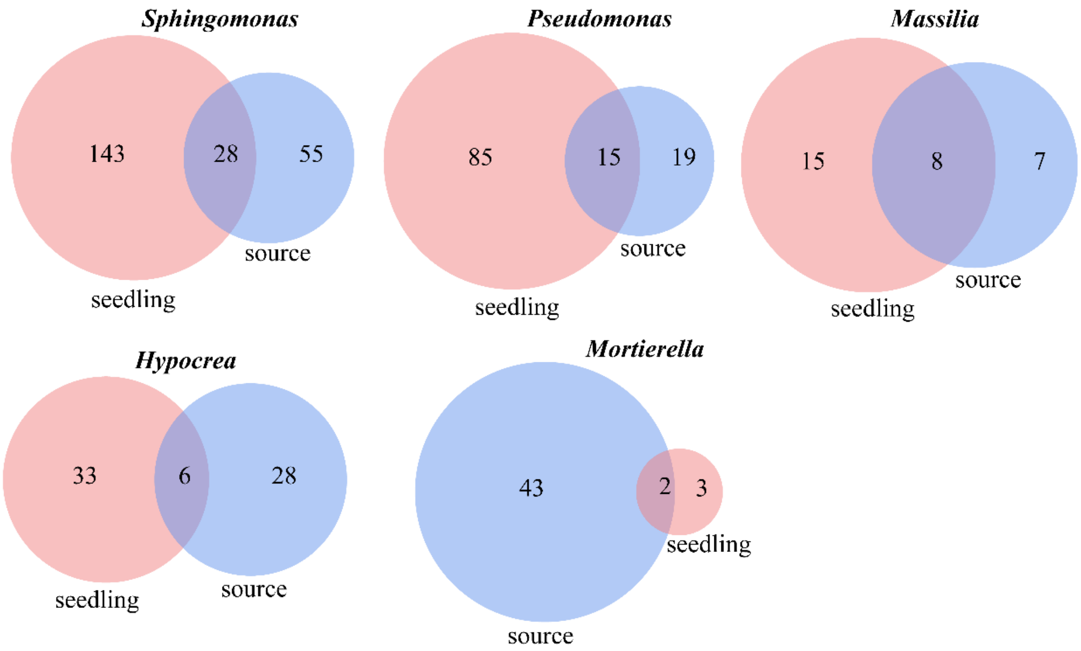


**Figure S8. Number of shared ASVs between the seedlings and inoculum sources (soil and litter) of several genera significantly correlated with seedling survival and growth.** Three bacterial genera (upper) and two fungal genera (bottom).

### 3. Supplementary tables

**Table S1 Potential functions of the core bacteria in *A. adenophora* rhizosphere soils.**

The bold text shows the N circle-related functional groups.

| **Function group** | **Relative abundance** |
|---|---|
| **nitrate reduction** | 0.0601 |
| **nitrate respiration** | 0.0594 |
| **nitrogen respiration** | 0.0594 |
| **nitrate denitrification** | 0.0590 |
| **nitrite denitrification** | 0.0590 |
| **nitrous oxide denitrification** | 0.0590 |
| **denitrification** | 0.0590 |
| **nitrite respiration** | 0.0590 |
| **nitrogen fixation** | 0.0326 |
| **ureolysis** | 0.0036 |
| chemoheterotrophy | 0.1432 |
| aerobic chemoheterotrophy | 0.1410 |
| photoheterotrophy | 0.0387 |
| phototrophy | 0.0387 |
| anoxygenic photoautotrophy S oxidizing | 0.0384 |
| anoxygenic photoautotrophy | 0.0384 |
| photoautotrophy | 0.0384 |
| predatory or exoparasitic | 0.0031 |
| aromatic compound degradation | 0.0027 |
| chitinolysis | 0.0021 |
| arsenate detoxification | 0.0008 |
| dissimilatory arsenate reduction | 0.0008 |
| plastic degradation | 0.0008 |
| fermentation | 0.0005 |
| dark oxidation of sulfur compounds | 0.0005 |
| invertebrate parasites | 0.0004 |
| human pathogens pneumonia | 0.0004 |
| human pathogens all | 0.0004 |
| human associated | 0.0004 |
| animal parasites or symbionts | 0.0004 |

**Table S2 Potential functions of the core bacteria in *A. adenophora* leaf litter.** The bold text shows the N circle-related functional groups.

| Function group | Relative abundance |
|---|---|
| **ureolysis** | 0.1568 |
| **nitrate respiration** | 0.0003 |
| **nitrate reduction** | 0.0003 |
| **nitrogen respiration** | 0.0003 |
| aerobic chemoheterotrophy | 0.3930 |
| chemoheterotrophy | 0.3930 |
| plant pathogen | 0.0229 |
| human pathogens all | 0.0108 |
| human associated | 0.0108 |
| animal parasites or symbionts | 0.0108 |
| chitinolysis | 0.0003 |
| aromatic compound degradation | 0.0003 |
| photoheterotrophy | 0.0002 |
| phototrophy | 0.0002 |

**Table S3 Potential functional guilds of core fungi in *A. adenophora* rhizosphere soils.** The bold text shows plant pathogen-related guilds.

| Guild | Relative abundance |
|---|---|
| **Plant Pathogen** | 0.1909 |
| **Endophyte–Plant Pathogen** | 0.0385 |
| **Animal Pathogen-Endophyte-Fungal Parasite-Lichen Parasite-Plant Pathogen-Wood Saprotroph** | 0.0373 |
| **Animal Pathogen-Endophyte-Lichen Parasite-Plant Pathogen-Soil Saprotroph-Wood Saprotroph** | 0.0216 |
| **Animal Pathogen-Endophyte–Plant Pathogen-Undefined Saprotroph** | 0.0195 |
| **Endophyte–Plant Pathogen-Wood Saprotroph** | 0.0124 |
| **Animal Pathogen-Plant Pathogen-Undefined Saprotroph** | 0.0106 |
| **Endophyte-Lichen Parasite-Plant Pathogen-Undefined Saprotroph** | 0.0044 |
| **Plant Pathogen-Wood Saprotroph** | 0.0029 |
| **Animal Endosymbiont-Animal Pathogen-Endophyte–Plant Pathogen-Undefined Saprotroph** | 0.0011 |
| Algal Parasite-Bryophyte Parasite-Fungal Parasite-Undefined Saprotroph | 0.0069 |
| Animal Pathogen | 0.0508 |
| Animal Pathogen-Endophyte-Epiphyte-Undefined Saprotroph | 0.0159 |
| Animal Pathogen-Fungal Parasite-Undefined Saprotroph | 0.0712 |

| | |
|---|---|
| Bryophyte Parasite-Dung Saprotroph-Ectomycorrhizal-Fungal Parasite-Leaf Saprotroph-Plant Parasite-Undefined Saprotroph-Wood Saprotroph | 0.0076 |
| Dung Saprotroph-Undefined Saprotroph-Wood Saprotroph | 0.0034 |
| Ectomycorrhizal | 0.1633 |
| Endophyte-Insect Pathogen | 0.0121 |
| Endophyte-Litter Saprotroph-Soil Saprotroph-Undefined Saprotroph | 0.1730 |
| Fungal Parasite | 0.0009 |
| Insect Pathogen | 0.0066 |
| Leaf Saprotroph | 0.0016 |
| Undefined Saprotroph | 0.0108 |
| unassigned | 0.1366 |

**Table S4 Potential functional guilds of the core fungi in *A. adenophora* litter.** The bold text shows plant pathogen-related guilds.

| **Guild** | **Relative abundance** |
|---|---|
| **Animal Pathogen-Plant Pathogen-Undefined Saprotroph** | 0.4682 |
| **Animal Pathogen-Endophyte–Plant Pathogen-Wood Saprotroph** | 0.1971 |
| **Fungal Parasite-Plant Pathogen-Plant Saprotroph** | 0.0056 |
| **Endophyte–Plant Pathogen-Undefined Saprotroph** | 0.0045 |
| **Animal Pathogen-Endophyte-Ericoid Mycorrhizal-Plant Pathogen-Wood Saprotroph** | 0.0024 |
| **Plant Pathogen-Wood Saprotroph** | 0.0021 |
| **Plant Pathogen** | 0.0013 |
| **Animal Endosymbiont-Animal Pathogen-Endophyte–Plant Pathogen-Undefined Saprotroph** | 0.0008 |
| Animal Pathogen | 0.0109 |
| Animal Pathogen-Endophyte-Endosymbiont-Epiphyte-Soil Saprotroph-Undefined Saprotroph | 0.0113 |
| Animal Pathogen-Endophyte-Epiphyte-Undefined Saprotroph | 0.0790 |
| Dung Saprotroph-Nematophagous | 0.0008 |
| Fungal Parasite-Litter Saprotroph | 0.0094 |
| Fungal Parasite-Soil Saprotroph | 0.0023 |
| Undefined Saprotroph | 0.0706 |
| Undefined Saprotroph-Wood Saprotroph | 0.0023 |
| unassigned | 0.1312 |

**Table S5 Taxonomic information of 33 representative strains isolated from 40 dead seedlings.** The numbers in brackets represent the similarity between our sequences and the sequences in the UNITE database.

| Strain code | Number of isolates | Relative abundance | Taxonomy | accession numbers |
|---|---|---|---|---|
| Z25 | 8 | 0.0417 | k__Fungi(100);p__Ascomycota(100);c__Dothideomycetes(100);o__Pleosporales(100);f__Didymellaceae(100);g__Allophoma(90);s__Allophoma_cylindrispora(90);Allophoma_cylindrispora(90); | OR473401 |
| Z18 | 1 | 0.0052 | k__Fungi(100);p__Ascomycota(100);c__Dothideomycetes(100);o__Pleosporales(100);f__Didymellaceae(100);g__Allophoma(90);s__Allophoma_cylindrispora(90);Allophoma_cylindrispora(90); | OR473394 |
| Z26 | 1 | 0.0052 | k__Fungi(100);p__Ascomycota(100);c__Dothideomycetes(100);o__Pleosporales(100);f__Didymellaceae(100);g__Allophoma(90);s__Allophoma_cylindrispora(90);Allophoma_cylindrispora(90); | OR473402 |
| Z38 | 1 | 0.0052 | k__Fungi(100);p__Ascomycota(100);c__Sordariomycetes(100);o__Glomerellales(100);f__Plectosphaerellaceae(100);g__Plectosphaerella(100);s__Plectosphaerella_cucumerina(95);Plectosphaerella_cucumerina(95); | OR473411 |
| Z28 | 7 | 0.0365 | k__Fungi(100);p__Ascomycota(100);c__Dothideomycetes(100);o__Pleosporales(100);f__Pleosporaceae(100);g__Alternaria(100);unclassified;unclassified; | OR473404 |
| Z29 | 8 | 0.0417 | k__Fungi(100);p__Ascomycota(100);c__Dothideomycetes(100);o__Pleosporales(100);f__Didymellaceae(100);g__Stagonosporopsis(92);unclassified;unclassified; | OR473405 |
| Z31 | 1 | 0.0052 | k__Fungi(100);p__Ascomycota(100);c__Dothideomycetes(100);o__Pleosporales(100);f__Didymellaceae(100);g__Epicoccum(100);s__Epicoccum_nigrum(93);Epicoccum_nigrum(93); | OR473406 |

| | | | | |
|---|---|---|---|---|
| Z32 | 9 | 0.0469 | k__Fungi(100);p__Ascomycota(100);c__Dothideomycetes(100);o__Pleosporales(100);f__Pleosporaceae(100);g__Alternaria(100);unclassified;unclassified; | OR473407 |
| Z33 | 14 | 0.0729 | k__Fungi(100);p__Ascomycota(100);c__Dothideomycetes(100);o__Pleosporales(100);f__Pleosporaceae(100);g__Alternaria(100);unclassified;unclassified; | OR473408 |
| Z34 | 10 | 0.0521 | k__Fungi(100);p__Ascomycota(100);c__Dothideomycetes(100);o__Pleosporales(100);f__Pleosporaceae(100);g__Alternaria(100);unclassified;unclassified; | OR473409 |
| Z36 | 3 | 0.0156 | k__Fungi(100);p__Ascomycota(100);c__Dothideomycetes(100);o__Pleosporales(100);f__Didymellaceae(100);g__Boeremia(100);s__Boeremia_exigua(100);Boeremia_exigua(100); | OR473410 |
| Z44 | 5 | 0.0260 | k__Fungi(100);p__Ascomycota(100);c__Dothideomycetes(100);o__Pleosporales(100);f__Didymellaceae(100);g__Allophoma(98);unclassified;unclassified; | OR473417 |
| Z39 | 1 | 0.0052 | k__Fungi(100);p__Ascomycota(100);c__Dothideomycetes(100);o__Capnodiales(100);f__Cladosporiaceae(100);g__Cladosporium(100);unclassified;unclassified; | OR473412 |
| Z40 | 1 | 0.0052 | k__Fungi(100);p__Ascomycota(100);c__Dothideomycetes(100);o__Pleosporales(100);f__Didymellaceae(100);g__Didymella(99);unclassified;unclassified; | OR473413 |
| Z41 | 1 | 0.0052 | k__Fungi(100);p__Ascomycota(100);c__Dothideomycetes(100);o__Pleosporales(100);f__Didymellaceae(100);g__Didymella(99);unclassified;unclassified; | OR473414 |
| Z42 | 2 | 0.0104 | k__Fungi(100);p__Ascomycota(100);c__Dothideomycetes(100);o__Pleosporales(100);f__Didymellaceae(100);g__Allophoma(96);s__Allophoma_cylindrispora(96);Allophoma_cylindrispora(96); | OR473415 |
| Z43 | 3 | 0.0156 | k__Fungi(100);p__Ascomycota(100);c__Dothideomycetes(100);o__Pleosporales(100);f__Didymellaceae(100);g__Allophoma(96);s__Allophoma_cylindrispora(96);Allophoma_cylindrispora(96); | OR473416 |
| Z10 | 2 | 0.0104 | k__Fungi(100);p__Ascomycota(100);c__Dothideomycetes(100);o__Pleosporales(100);f__Didymellaceae(100);g__Epicoccum(100);unclassified;unclassified; | OR473386 |

| | | | | |
|---|---|---|---|---|
| Z46 | 3 | 0.0156 | k__Fungi(100);p__Ascomycota(100);c__Sordariomycetes(100);o__Glomerellales(100);f__Glomerellaceae(100);g__Colletotrichum(100);s__Colletotrichum_clavatum(100);Colletotrichum_clavatum(100); | OR473418 |
| Z27 | 77 | 0.4010 | k__Fungi(100);p__Ascomycota(100);c__Dothideomycetes(100);o__Pleosporales(100);f__Didymellaceae(100);g__Allophoma(90);s__Allophoma_cylindrispora(90);Allophoma_cylindrispora(90); | OR473403 |
| Z24 | 5 | 0.0260 | k__Fungi(100);p__Ascomycota(100);c__Dothideomycetes(100);o__Pleosporales(100);f__Didymellaceae(99);g__Didymella(99);unclassified;unclassified; | OR473400 |
| Z23 | 8 | 0.0417 | k__Fungi(100);p__Ascomycota(100);c__Dothideomycetes(100);o__Pleosporales(100);f__Pleosporaceae(100);g__Alternaria(100);unclassified;unclassified; | OR473399 |
| Z22 | 2 | 0.0104 | k__Fungi(100);p__Ascomycota(100);c__Sordariomycetes(100);o__Hypocreales(100);f__Nectriaceae(100);g__Fusarium(99);unclassified;unclassified; | OR473398 |
| Z21 | 4 | 0.0208 | k__Fungi(100);p__Ascomycota(100);c__Dothideomycetes(100);o__Pleosporales(100);f__Didymellaceae(100);g__Epicoccum(100);unclassified;unclassified; | OR473397 |
| Z20 | 3 | 0.0156 | k__Fungi(100);p__Ascomycota(100);c__Dothideomycetes(100);o__Pleosporales(99);f__Didymellaceae(99);g__Epicoccum(99);s__Epicoccum_nigrum(91);Epicoccum_nigrum(91); | OR473396 |
| Z19 | 3 | 0.0156 | k__Fungi(100);p__Ascomycota(100);c__Sordariomycetes(100);o__Hypocreales(100);f__Nectriaceae(100);g__Fusarium(100);s__Fusarium_kyushuense(100);Fusarium_kyushuense(100); | OR473395 |
| Z17 | 1 | 0.0052 | k__Fungi(100);p__Ascomycota(100);c__Dothideomycetes(100);o__Capnodiales(100);f__Cladosporiaceae(100);g__Cladosporium(100);unclassified;unclassified; | OR473393 |
| Z16 | 1 | 0.0052 | k__Fungi(100);p__Ascomycota(100);c__Dothideomycetes(100);o__Pleosporales(100);f__Didymellaceae(100);g__Stagonosporopsis(100);s__Stagonosporopsis_cucurbitacearum(93);Stagonosporopsis_cucurbitacearum(93); | OR473392 |

| | | | | |
|---|---|---|---|---|
| Z15 | 1 | 0.0052 | k__Fungi(100);p__Ascomycota(100);c__Dothideomycetes(100);o__Pleosporales(100);f__Pleosporaceae(100);g__Alternaria(100);s__Alternaria_destruens(100);Alternaria_destruens(100); | OR473391 |
| Z14 | 1 | 0.0052 | k__Fungi(100);p__Ascomycota(100);c__Dothideomycetes(100);o__Pleosporales(100);f__Pleosporaceae(100);g__Alternaria(100);s__Alternaria_destruens(97);Alternaria_destruens(97); | OR473390 |
| Z13 | 1 | 0.0052 | k__Fungi(100);p__Ascomycota(100);c__Dothideomycetes(100);o__Pleosporales(98);f__Didymellaceae(98);g__Epicoccum(98);s__Epicoccum_nigrum(91);Epicoccum_nigrum(91); | OR473389 |
| Z12 | 2 | 0.0104 | k__Fungi(100);p__Ascomycota(100);c__Sordariomycetes(100);o__Sordariales(100);f__Chaetomiaceae(100);g__Chaetomium(100);unclassified;unclassified; | OR473388 |
| Z11 | 2 | 0.0104 | k__Fungi(100);p__Ascomycota(100);c__Sordariomycetes(100);o__Hypocreales(100);f__Nectriaceae(100);g__Fusarium(99);unclassified;unclassified; | OR473387 |

**Table S6 PERMANOVA of bacterial communities and function at each inoculation time.** Bold shows $P < 0.05$.

| Inoculation time | Factor | Df | Bacterial community | | | Bacterial function | | |
|---|---|---|---|---|---|---|---|---|
| | | | $R^2$ | F.Model | Pr(>F) | $R^2$ | F.Model | Pr(>F) |
| G0 | compartment | 1 | 0.132 | 1.761 | **0.045** | 0.133 | 1.423 | 0.272 |
| | nutrition | 1 | 0.194 | 2.582 | **0.004** | 0.023 | 0.249 | 0.8 |
| | source | - | - | - | - | - | - | - |
| G21 | compartment | 1 | 0.145 | 4.672 | **0.001** | 0.197 | 6.699 | **0.005** |
| | nutrition | 1 | 0.067 | 2.151 | **0.031** | 0.037 | 1.249 | 0.276 |
| | source | 1 | 0.167 | 5.384 | **0.001** | 0.180 | 6.137 | **0.002** |
| G21+28 | compartment | 1 | 0.200 | 5.987 | **0.001** | 0.144 | 3.977 | **0.026** |
| | nutrition | 1 | 0.062 | 1.839 | 0.085 | 0.019 | 0.516 | 0.622 |
| | source | 1 | 0.069 | 2.056 | **0.041** | 0.197 | 6.699 | **0.005** |
| G28 | compartment | 1 | 0.174 | 5.391 | **0.001** | 0.228 | 8.444 | **0.001** |
| | nutrition | 1 | 0.078 | 2.419 | **0.012** | 0.061 | 2.262 | 0.092 |
| | source | 1 | 0.102 | 3.148 | **0.002** | 0.170 | 6.288 | **0.001** |

**Table S7 PERMANOVA of fungal communities and function at each inoculation time.** Bold shows $P < 0.05$.

| Inoculation time | Factor | Df | Fungal community | | | Fungal function | | |
|---|---|---|---|---|---|---|---|---|
| | | | $R^2$ | F.Model | Pr(>F) | $R^2$ | F.Model | Pr(>F) |
| G0 | compartment | 1 | 0.059 | 0.656 | 0.935 | 0.059 | 0.614 | 0.872 |
| | nutrition | 1 | 0.128 | 1.414 | 0.092 | 0.071 | 0.740 | 0.755 |
| | source | - | - | - | - | - | - | - |
| G21 | compartment | 1 | 0.059 | 1.515 | 0.101 | 0.117 | 3.214 | **0.013** |
| | nutrition | 1 | 0.048 | 1.223 | 0.213 | 0.016 | 0.449 | 0.917 |
| | source | 1 | 0.110 | 2.797 | **0.006** | 0.140 | 3.867 | **0.01** |
| G21+28 | compartment | 1 | 0.091 | 2.380 | **0.034** | 0.126 | 3.714 | **0.011** |
| | nutrition | 1 | 0.049 | 1.273 | 0.161 | 0.062 | 1.829 | 0.095 |
| | source | 1 | 0.091 | 2.372 | **0.025** | 0.136 | 4.033 | **0.006** |
| G28 | compartment | 1 | 0.061 | 1.539 | 0.099 | 0.111 | 3.228 | **0.011** |
| | nutrition | 1 | 0.041 | 1.047 | 0.358 | 0.049 | 1.418 | 0.207 |
| | source | 1 | 0.108 | 2.724 | **0.007** | 0.154 | 4.503 | **0.001** |

**Table S8 Information about the abbreviations for the fungal functional guilds shown in Fig. 6D-E.**

| Fungal guild guild shown in Fig. 6D-E | Abbreviation of fungal function guild |
|---|---|
| Dung Saprotroph-Nematophagous | Dung Saprotroph-N |
| Dung Saprotroph-Plant Saprotroph-Soil Saprotroph | Dung Saprotroph-PSSS |
| Plant Pathogen-Undefined Parasite-Undefined Saprotroph | Plant Pathogen-UPUS |
| Ectomycorrhizal-Undefined Saprotroph | Ectomycorrhizal-US |
| Endophyte–Plant Pathogen | Endophyte-PP |
| Ectomycorrhizal-Fungal Parasite-Soil Saprotroph-Undefined Saprotroph | Ectomycorrhizal-FPSSUS |
| Litter Saprotroph-Plant Pathogen | Litter Saprotroph-PP |
| Ectomycorrhizal-Fungal Parasite-Leaf Saprotroph-Plant Parasite-Undefined Saprotroph-Wood Saprotroph | Ectomycorrhizal-FPLSPPUSWS |
| Endophyte-Litter Saprotroph-Soil Saprotroph-Undefined Saprotroph | Endophyte-LSSSUS |
| Fungal Parasite-Plant Pathogen-Plant Saprotroph | Fungal Parasite-PPPS |
| Animal Endosymbiont-Animal Pathogen-Endophyte–Plant Pathogen-Undefined Saprotroph | Animal Endosymbiont-APEPPUS |
| Dung Saprotroph-Ectomycorrhizal-Litter Saprotroph-Undefined Saprotroph | Dung Saprotroph-ELSUS |
| Animal Pathogen-Endophyte-Ericoid Mycorrhizal-Plant Pathogen-Wood Saprotroph | Animal Pathogen-EEMPPWS |
| Dung Saprotroph-Wood Saprotroph | Dung Saprotroph-WS |
| Endophyte–Plant Pathogen-Undefined Saprotroph | Endophyte-PPUS |

**Table S9 Fertilizer content in the Pindstrup substrate.**

| | **Content(g/$m^3$)** |
|---|---|
| Nitrate-N | 4.536 |
| Ammonium-N | 3.24 |
| Phosphorus (P) | 3.888 |
| Phosphorus (P2O5) | 9.072 |
| Potassium (K) | 12.96 |
| Potassium (K20) | 15.552 |
| Magnesium (Mg) | 0.9072 |
| Magnesium (MgO) | 1.4904 |
| Boron (B) | 0.02592 |
| Molybdenum (Mo) | 0.1296 |
| Copper (Cu) | 0.11016 |
| Manganese (Mn) | 0.18792 |
| Zinc (Zn) | 0.05832 |
| Iron (Fe) | 0.54432 |
| Wetting agent | 100 (mL/$m^3$) |
| pH | 6 |
| EC (ms/m) | 40 |

**Note:** All fertilizer content data for the Pindstrup substrate were obtained from https://www.pindstrup.com.

### References


Drummond, A. J., Suchard, M. A., Xie, D., & Rambaut, A. (2012). Bayesian Phylogenetics with BEAUti and the BEAST 1.7. *Molecular Biology and Evolution*, *29*(8), 1969-1973. https://doi.org/10.1093/molbev/mss075

Edgar, R. C. (2004). MUSCLE: multiple sequence alignment with high accuracy and high throughput. *Nucleic Acids Research*, *32*(5), 1792-1797. https://doi.org/10.1093/nar/gkh340

Nilsson, R. H., Larsson, K.-H., Taylor, A. F S., Bengtsson-Palme, J., Jeppesen, T. S., Schigel, D., Kennedy, P., Picard, K., Glöckner, F. O., Tedersoo, L., Saar, I., Kõljalg, U., & Abarenkov, K. (2018). The UNITE database for molecular identification of fungi: handling dark taxa and parallel taxonomic classifications. *Nucleic Acids Research*, *47*(D1), D259-D264. https://doi.org/10.1093/nar/gky1022.

Quast, C., Pruesse, E., Yilmaz, P., Gerken, J., Schweer, T., Yarza, P., Peplies, J., & Glockner, F. O. (2013). The SILVA ribosomal RNA gene database project: improved data processing and web-based tools. *Nucleic Acids Research*, *41*(Database issue), D590-596. https://doi.org/10.1093/nar/gks1219

Stewart, C. N., & Via, L. E. (1993). A rapid CTAB DNA isolation technique useful for rapid fingerprinting and other PCR applications. *Biotechniques*, *14*(5), 748-750. <Go to

ISI>://WOS:A1993LA81200016

Tamura, K., Stecher, G., Peterson, D., Filipski, A., & Kumar, S. (2013). MEGA6: Molecular Evolutionary Genetics Analysis Version 6.0. *Molecular Biology and Evolution*, *30*(12), 2725-2729. https://doi.org/10.1093/molbev/mst197

Zaret, M. M., Bauer, J. T., Clay, K., & Whitaker, B. K. (2021). Conspecific leaf litter induces negative feedbacks in Asteraceae seedlings. *Ecology*, *102*(12), e03557. https://doi.org/10.1002/ecy.3557

Zhang, C., Willis, C. G., Burghardt, L. T., Qi, W., Liu, K., de Moura Souza-Filho, P. R., Ma, Z., & Du, G. (2014). The community-level effect of light on germination timing in relation to seed mass: a source of regeneration niche differentiation. *New Phytol*, *204*(3), 496-506. https://doi.org/https://doi.org/10.1111/nph.12955
